# Pupil-associated states modulate excitability but not stimulus selectivity in primary auditory cortex

**DOI:** 10.1101/619619

**Authors:** Zachary P. Schwartz, Brad N. Buran, Stephen V. David

## Abstract

Recent research in mice indicates that luminance-independent fluctuations in pupil size predict variability in spontaneous and evoked activity of single neurons in auditory and visual cortex. These findings suggest that pupil is an indicator of large-scale changes in arousal state that affect sensory processing. However, it is not known whether pupil-related state also influences the selectivity of auditory neurons. We recorded pupil size and single-unit spiking activity in the primary auditory cortex (A1) of non-anesthetized male and female ferrets during presentation of natural vocalizations and tone stimuli that allow measurement of frequency and level tuning. Neurons showed a systematic increase in both spontaneous and sound-evoked activity when pupil was large, as well as desynchronization and a decrease in trial-to-trial variability. Relationships between pupil size and firing rate were non-monotonic in some cells. In most neurons, several measurements of tuning, including acoustic threshold, spectral bandwidth, and best frequency, remained stable across large changes in pupil size. Across the population, however, there was a small but significant decrease in acoustic threshold when pupil was dilated. In some recordings, we observed rapid, saccade-like eye movements during sustained pupil constriction, which may indicate sleep. Including the presence of this state as a separate variable in a regression model of neural variability accounted for some, but not all, of the variability and non-monotonicity associated with changes in pupil size.

**New & Noteworthy:** Cortical neurons vary in their response to repeated stimuli, and some portion of the variability is due to fluctuations in network state. By simultaneously recording pupil and single-neuron activity in auditory cortex of ferrets, we provide new evidence that network state affects the excitability of auditory neurons, but not sensory selectivity. In addition, we report the occurrence of possible sleep states, adding to evidence that pupil provides an index of both sleep and physiological arousal.

## Introduction

Although the iris primarily functions to regulate the formation of images on the retina, pupil dilation and constriction are also a result of autonomic nervous activity unrelated to visual stimuli (Loewenfeld and Lowenstein, 1999). Growing evidence suggests that these autonomic changes in pupil diameter are an indicator of changes in internal state that affect processing in sensory cortex (Reimer et al., 2014; McGinley et al., 2015a; Vinck et al., 2015). In mice, pupil dilation is correlated with transitions from stillness to walking (McGinley et al., 2015a; Vinck et al., 2015), which increase firing rates in visual cortex (Niell and Stryker, 2010) and decrease firing rates in auditory cortex (Schneider et al., 2014). Dilated pupil is associated with increases in high-frequency local field potential (LFP), a physiological signature of alertness, in sensory cortex (Harris and Thiele, 2011; Vinck et al., 2015). In mouse auditory cortex, intermediate pupil diameter is also associated with increases in the gain of sound-evoked responses, decreased spontaneous activity, and stable, hyperpolarized subthreshold membrane potential, which have been hypothesized to reflect an “optimal state” for detecting sensory stimuli (McGinley et al., 2015a).

Pupil dilation has also been correlated with changes in sensory selectivity. In visual cortex, orientation tuning becomes more selective when pupil dilates, suggesting that sensory representations are more precise in some internal brain states (Reimer et al., 2014). However, it is not known if pupil-associated state influences the selectivity of neurons in the auditory system. Some population-level associations between pupil size and cortical activity do differ between sensory systems (Shimaoka et al., 2018). Data from mouse auditory cortex – which suggests that pupil size is non-monotonically related to neural activity (McGinley et al., 2015a) – conflicts with data from mouse visual cortex, which show either monotonic increases in neural activity when pupil dilates (Reimer et al., 2014; Vinck et al., 2015), or simultaneous increases and decreases in different neurons recorded at the same time (Stringer et al., 2019). These observations may reflect functional differences between the auditory and visual systems.

To explore effects of pupil-associated state on sensory coding in auditory cortex, we recorded pupil size and single-unit spiking activity in the primary auditory cortex (A1) of head-restrained, non-anesthetized ferrets. To study effects of internal state on gain and spontaneous activity, we presented a large number of repetitions (up to 120) of ferret vocalizations. To study state effects on sensory selectivity we presented random sequences of tone pips at various frequencies and levels. Each stimulus set therefore allowed in-depth exploration of a distinct feature of auditory responses, as well as comparison to previous work in the visual system (Reimer et al., 2014). We characterized effects of pupil by comparing responses to each distinct sound as pupil size varied. Neurons typically showed an increase in baseline firing rate as well as the gain and reliability of sound-evoked activity when pupil was large, although there was also evidence for non-monotonic effects of pupil-associated state in some neurons. In most neurons, parametric measurements of tuning, including acoustic threshold, spectral bandwidth, and best frequency, remained stable across large changes in pupil size. Across the population, however, we observed a small decrease in acoustic threshold when pupil was dilated.

In some recordings, we observed increased saccadic eye movements during periods of sustained pupil constriction, possibly indicating sleep onset. Including the presence of this state as a separate variable in a regression model of neural variability accounted for some, but not all, of the variability associated with changes in pupil size. The stability of best frequency across large changes in pupil size (and associated changes in gain and spontaneous activity) contrasts with task-related plasticity of receptive fields in ferret auditory cortex (Fritz et al., 2003; David et al., 2012), suggesting a difference between effects of engaging in tasks that require listening and variation within passive states.

## Methods

All procedures were approved by the Oregon Health and Science University Institutional Animal Care and Use Committee (IACUC) and conform to standards of the Association for Assessment and Accreditation of Laboratory Animal Care (AAALAC).

### Neurophysiology

Five young adult ferrets (4 males and 1 female, Marshall Farms) were implanted with stainless steel headpost and acrylic cap to permit head-fixation and access to auditory cortex. After recovery (1-2 weeks), animals were habituated to a head-fixed posture and presentation of auditory stimuli. Following habituation, a small (1-2 mm) craniotomy was opened over primary auditory cortex (A1). In 66 recordings, data was collected using 1-4 independently positioned tungsten microelectrodes (FHC, 1-5 MΩ) advanced by independent microdrives (Electrode Positioning System, Alpha-Omega). In 3 recordings, data was collected using a single-shank 64-channel silicon microelectrode array, spanning 1 mm (Masmanidis Lab, UCLA) (Du et al., 2011).

After a unit was isolated during tungsten recordings, its auditory responsiveness was tested using tones or narrowband noise bursts. If the site showed no response, the electrode was lowered to search for another site. During depth-array recordings, the probe was lowered into auditory cortex until auditory-evoked activity was observed across the span of recorded channels. Amplified and digitized signals were stored using open-source data acquisition software (Englitz et al., 2013; Siegle et al., 2017). Ferrets were observed by video monitor during recordings. If a unit became indistinguishable from the noise floor following a body movement, the recording was stopped and the 5 trials preceding the putative isolation loss (28 to 40 seconds of recording time) were excluded from analysis. A1 recording sites were identified by characteristic short latency, frequency selectivity, and tonotopic gradients of single or multiunit activity across multiple penetrations (Bizley et al., 2005; Elgueda et al., 2019).

### Spike sorting

For tungsten electrode recordings, events were extracted from the continuous electrophysiological signal using a principal components analysis and k-means clustering (Meska-PCA, NSL). Single-unit isolation was quantified from cluster overlap as the fraction of spikes likely to be produced by a single neuron rather than another unit. For depth-array recordings, single-unit spikes were sorted using polytrode sorting software (Pachitariu et al., 2016). Only units maintaining stable isolation > 95% through the experiment were considered single neurons for analysis.

### Pupillometry

During neurophysiological recordings, infrared video of one eye was collected for offline measurement of pupil size. Video was collected using either (1) a commercial CCTV camera fitted with a lens to provide magnification of the eye and a band-stop infrared filter (Heliopan ES 49 RG 780) or (2) an open-source camera (Adafruit TTL Serial Camera) fitted with a lens (M12 Lenses PT-2514BMP 25.0 mm) whose focal length allowed placement of the camera 10 cm from the ferret’s eye. To improve contrast, the imaged eye was illuminated by a bank of infrared LEDs. Ambient luminance was provided using a ring light (AmScope LED-144S). At the start of each recording day, the intensity of the ring light was set to a level (∼1500 lux measured at the recorded eye) chosen to give a maximum dynamic range of pupil sizes. Light intensity remained fixed across the recording session.

Pupil size was measured using custom MATLAB (version R2016b) software (code available at https://bitbucket.org/lbhb/baphy). For each video, an intensity threshold was selected to capture pupil pixels and exclude the surrounding image. During initial recordings, the threshold was selected manually. We observed that the intensity histogram is multi-modal with the first peak (reflecting the darkest pixels) generally corresponding to the pupil. In later recordings, we therefore automatically updated the threshold of each frame to position it at the first valley in the frame’s intensity histogram. Each frame was smoothed by a Gaussian filter before thresholding, then segmented by Moore boundary tracing. The segment with the largest area was identified as pupil. We measured pupil size as the length of the minor axis of an ellipse fit to this region using a direct least-squares method. To avoid identifying shadows at the edge of the eye or other dark regions of the image as pupil, we restricted the search for the largest-area segment to a rectangular region of interest surrounding the pupil identified in the preceding frame (Nguyen and Stark, 1993). Measurements were calibrated by comparing the width of each ferret’s eye (in mm) and the width of the ferret eye as it appeared in each video (in pixels), which gave a conversion factor of pixels to mm.

The frame rate of the cameras used in the experiments varied from 10 to 30 frames/second. To compensate for variability in camera frame rate, we recorded a timestamp at the start and end of each trial, then interpolated measurements of pupil size to match the sampling of the simultaneously recorded neural data. This procedure ensured that the two data streams (video and neural recording) remained synchronized throughout each recording.

Blink artifacts were identified by rapid, transient changes in pupil size (McGinley et al., 2015a). The derivative of the pupil trace was taken and bins with derivatives more than 6 standard deviations from the mean were marked. Blinks were identified within these bins by screening for decreases in pupil size followed by increases. Data during a 1-second period surrounding the blink was then removed from the trace and replaced by a linear interpolation of the pupil size immediately before and after the blink.

When comparing pupil and neural data, a 750 ms offset was applied to pupil trace to account for the lagged relationship between changes in pupil size and neural activity in auditory cortex to allow for comparison with previous research (McGinley et al., 2015a). Setting the offset at values between 0 and 1.5 s did not affect the mean accuracy of the pupil-based regression models of neural activity described below (data not shown).

To measure changes in eye position, we calculated the Euclidean distance between the center of the ellipse fit to the pupil in each video frame, then multiplied by frame rate to find eye speed. We identified putative sleep states by screening for a combination of tonically constricted pupil and a high rate of saccades. Sleep states were identified by selecting parameter values for each recording (maximum pupil size, maximum pupil standard deviation, minimum saccade speed, minimum saccade rate) that marked abrupt state transitions in the traces of pupil size and eye speed. Gaps of less than 15 seconds between sleep episodes were considered artifacts and coded as sleep. We restricted our analysis to sleep episodes with a duration of at least 30 seconds.

### Acoustic stimuli

Stimulus presentation was controlled by custom MATLAB software (code available at https://bitbucket.org/lbhb/baphy). Digital acoustic signals were transformed to analog (National Instruments), amplified (Crown), and delivered through a free-field speaker (Manger).

During initial recording sessions, ferrets were presented with two 3-second species-specific vocalizations: an infant call and an adult aggression call. Stimuli were recorded in a sound-attenuating chamber using a commercial digital recorder (44-kHz sampling, Tascam DR-400). No animal that produced the recorded vocalizations was used in the study. Each vocalization was repeated up to 120 times. The order of vocalizations was randomized on each repetition. Each repetition was preceded by a 2-second silent period to allow measurement of spontaneous neural activity on each trial.

To improve sampling of internal states, recording was paused after sound repetition 40 and 80 for several seconds, during which the ferret was roused by an unexpected sound (turning the doorknob of the recording booth), which often resulted in pupil dilation. A stepwise regression analysis (see below) using the number of trials since each pause to predict neural activity suggests that the time since this event did not predict neural activity independent of pupil size (data not shown).

During later recording sessions, ferrets were presented with sequences of 10-ms tone pips with varying frequency (4-6 octaves surrounding best frequency, ranging over 32 Hz to 40 kHz) and level (0 to 65 dB SPL, 5 or 10 dB SPL steps). Each pip sequence was 6 seconds long, preceded by a 2-second silent period to allow measurement of spontaneous neural activity. Tone-pip sequences were repeated multiple times in each recording (mean repetitions +/− SEM: 28 +/− 2). During some recordings, tone-pip sequences were interleaved with ferret vocalizations to improve sampling of pupil states in each data set.

### Data analysis: Gain and baseline firing rate

Neural and pupil data for ferret vocalization recordings was binned at 4 Hz. Peristimulus time histogram (PSTH) responses to each vocalization were calculated by aligning spike activity to stimulus onset and averaging across presentations. To isolate evoked activity, the mean spike rate during the two seconds of silence preceding stimulus onset was subtracted from the PSTH.

Neural and pupil data for responses to tone-pip sequences were binned at 100 Hz. A frequency response area (FRA) was calculated by taking the mean spike rate for each sound during a window 10 ms to 60 ms following pip onset. To isolate evoked activity, the mean spike rate during the two seconds of silence preceding tone-pip sequence onset was subtracted from the FRA.

We used multivariate regression models to examine effects of pupil state on spontaneous and sound-evoked activity. We modeled the response to each presentation of a tone pip or vocalization as a function of average spontaneous rate over the entire recording, average response to the stimulus across all trials, and pupil diameter during each stimulus presentation.

The *baseline-gain* model accounted for a linear relationship between pupil diameter and both spontaneous and evoked spike rate. According to this model, single-trial activity, *r*_i_(*t*, *s*), is

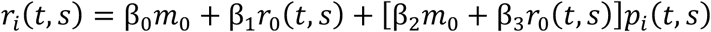

Here, *r*_0_(*t*, *s*) is the mean time-varying PSTH response evoked by stimulus *s* (either one vocalization or one tone pip at a single frequency and level), *m*_0_ is the mean spontaneous rate, and *p*_*i*_(*t*, *s*) is pupil size observed simultaneously to *r*_*i*_(*t*, *s*). The model coefficients indicate how pupil the neuron’s overall spiking activity (baseline, *β*_2_) and the scaling of the cell’s sound-evoked response (gain, *β*_3_) vary with pupil size. To cast all *β* parameters in units of pupil^-1^, we normalized the baseline terms by *m*_0_. We therefore refer to this as the “baseline-gain” or “first-order baseline-gain” model.

Unless otherwise specified, model parameters were fit by linear, least-squares regression using 20-fold cross-validation. The median model parameters across all 20 fits was used to compare coefficients across cells. Model accuracy for a given cell was quantified by the squared Pearson correlation between actual single-trial firing rates and firing rates predicted using the cross-validated model coefficients. Significance of model fits was assessed using a permutation test. A cell with a *significant fit* was one in which a fit to real pupil data showed accuracy greater than a model fit to pupil data shuffled across trials for at least 50 of 1000 shuffles (i.e., predictions improved with *p* < 0.05 of occurring by chance).

To determine what aspects of the baseline-gain model were important for the capturing the relationship between pupil and firing rate, we compared it to several variants. The *baseline only* model allowed only the baseline rate, but not the gain of the evoked response, to scale with pupil,

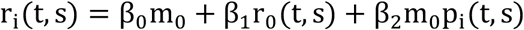

The *gain only* model allowed only the gain of the evoked response, and not the baseline rate, to scale with pupil,

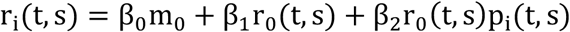

We also considered nonlinear relationships between pupil and spike rate. The *second-order baseline* model allowed the baseline rate to scale with the square of pupil size, allowing non-monotonic effects of internal state,

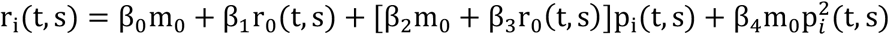

The *second-order baseline-gain* model allowed both the baseline rate and gain to scale with the square of pupil size,

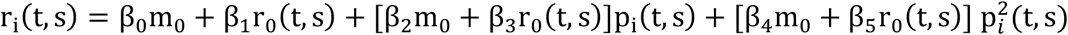

We measured the performance of each model by comparing its ability to predict single-trial data to the PSTH or FRA alone, without any information on pupil size (the *null model*),

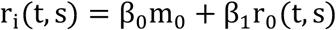

For each cell and each model, we measured the percent improvement in accuracy (i.e., the difference in squared Pearson correlation coefficient, *R*^2^, between the model predictions and actual data) over the null model,

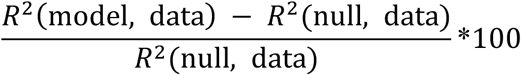

Each model’s performance was quantified as the median improvement in accuracy across all cells that showed a significant fit for any tested model (*p* < 0.05, permutation test). For comparing model performance across collections of neurons, we performed a Wilcoxon signed-rank test (*sign test*) between the distribution of prediction correlations across neurons for each model.

### Data analysis: Non-monotonicity

A previous report (McGinley et al., 2015a) observed non-monotonic relationships between pupil size and neural activity in mouse auditory cortex. Following its language, we refer to neurons with a maximum firing rate at intermediate pupil sizes and a lesser firing rate at small and large pupil sizes as “inverted U”s. We refer to neurons with a minimum firing rate at intermediate pupil sizes as “U”s.

We restricted our analysis of non-monotonicity to neurons that showed a significant fit for the second-order baseline-gain model relative to the same model fit to shuffled pupil data (*n* = 79/114 neurons). The second-order terms of this model allow for the possibility of a non-monotonic relationships between pupil size and firing rate.

We used a segmented linear model to test for non-monotonic relationships between pupil size and firing rate in single neurons (Simonsohn, Uri, 2018). The segmented model is similar to the baseline-gain model (see above), except that data is partitioned into intervals at a *breakpoint* in pupil size. The fit parameters of the model may differ on either side of the breakpoint. This allows a variety of relationships between pupil and firing rate, including linear increases and decreases, Us, and inverted Us.

The segmented model included separate breakpoints for pupil-related effects on baseline and gain (β_0_, β_4_). Both breakpoints were fit simultaneously with other free parameters in the model. Baseline and gain effects were combined to estimate the neuron’s firing rate, i.e.,

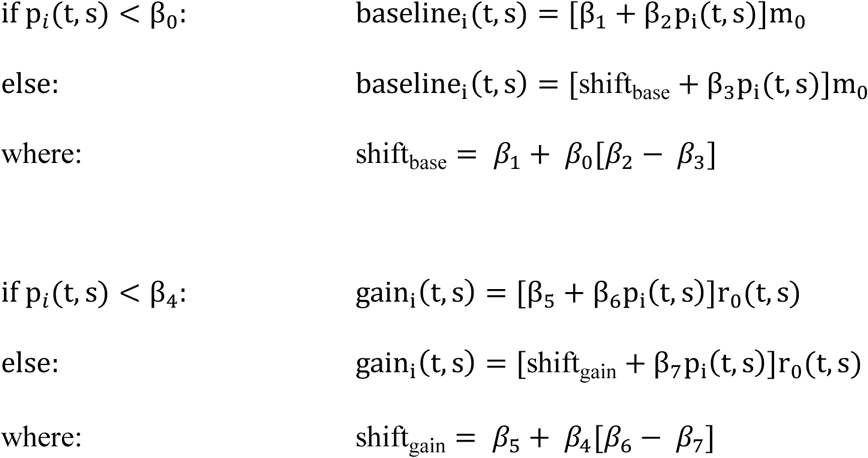

and

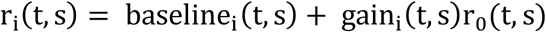

The constraints on the two *shift* parameters ensure that the functions will be continuous at their breakpoints. Model parameters were fit by nonlinear least-squares using the trust-region-reflective algorithm and 20-fold cross validation. A change in the sign of pupil coefficients across the breakpoint (*β*_2_ versus *β*_3_ or *β*_6_ versus *β*_7_) indicates a non-monotonic relationship between pupil and firing rate. The direction of the sign change indicates if a cell is a U or inverted U.

To identify non-monotonic neurons, we compared the accuracy of the segmented linear model to an identical model that was constrained to be monotonic. That is, the slope coefficients related to baseline (*β*_2_, *β*_3_) and gain (*β*_6_, *β*_7_) were constrained to have the same sign on both sides of the breakpoint. If the non-constrained model showed a sign change as well as an improvement in accuracy over the constrained model, we classified the cell as showing a *non-monotonic trend*.

To identify *significantly non-monotonic* neurons, we performed a permutation test. We randomly shuffled half the firing rate predictions between the constrained and non-constrained models, then calculated the difference in accuracy between the shuffled prediction rates. Significantly non-monotonic neurons were those in which less than 50 of 1000 shuffles showed a greater difference in accuracy than that observed between the constrained and non-constrained models (*p* < 0.05 of a chance improvement).

### Data analysis: Frequency and level tuning

To assess changes in frequency and level tuning associated with pupil state, neural responses to tone-pip sequences were binned at 100 Hz. Data from each tone-pip recording was divided into two bins (*large-pupil* and *small-pupil*) based on the median pupil size during the recording. Data from tone-pip sequences was analyzed using: (1) the 2-second silence preceding each tone-pip sequence (*spontaneous activity*), (2) the interval from 60 ms to 10 ms preceding onset of each tone pip (*prestimulus activity*), and (3) the interval 10 ms to 60 ms after the onset of each tone pip (*stimulus-evoked activity*). Since the FRA was assessed using a cloud of tone pips presented in rapid succession, the cell’s firing rate during the 50 ms preceding sound onset (i.e., prestimulus activity) was a better measure of the cell’s baseline response for sub-threshold levels than spontaneous activity.

For display, examples of FRAs were calculated taking the mean stimulus-evoked spike rate for each tone pip, smoothing using a two-dimensional box filter with dimensions equivalent to 15 dB and 1.5 octaves, and subtracting the mean spontaneous activity across all tone-pip sequences in the recording. To measure changes in spontaneous and driven rates in the tone-pip data without exploring changes in tuning, we used the mean spontaneous rate and the difference between the mean stimulus-evoked and prestimulus firing rates. Then to measure possible changes in sensory tuning we fit hierarchical regression models to the prestimulus and stimulus-evoked spike data for each tone pip.

The hierarchical models were implemented in STAN (http://mc-stan.org) and fit using Bayesian inference. In contrast to conventional model fitting, Bayesian analysis allows for simple calculations of credible intervals on derived parameters (e.g., parameters that are mathematical functions of fitted coefficients), is more robust to outliers through the use of Poisson likelihoods for spike count data, and offers simple construction of realistic hierarchical models (Gelman et al., 2003). Each model was fit four times for 1000 samples after a 1000 sample iteration burn-in period. Posterior samples were combined across all chains for inference.

### Data analysis: Rate-level functions

To estimate changes in level tuning linked to internal state, we first calculated characteristic frequency (CF) before binning data by pupil size. An FRA for all data from the recording was calculated using data from the stimulus-evoked epoch and smoothed with a two-dimensional box filter with dimensions equivalent to 15 dB and 1.5 octaves. A separate standard error was calculated for each coefficient of the FRA by measuring variance across repetitions of tone pips at that frequency and level. The minimum sound level that evoked a response at least 2 standard errors above the mean spike rate during the 100 ms preceding pip onset, measured across all pips, was calculated for each frequency. CF was defined as the frequency that required the minimum sound level to evoke a response.

For each cell, we went back to the unsmoothed FRA data, binned the tone-evoked responses according to pupil size and generated rate-level functions. Rate-level functions were smoothed by averaging the response across the three frequencies closest to CF. Rate-level functions for individual cells were fit using a function with three parameters, baseline firing rate, *b*, slope, *s*, and threshold, *θ*, for each pupil condition:

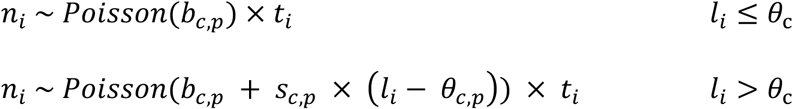

Here, *n* is the number of spikes observed in a time interval, *t*, for cell, *c*, and pupil condition, *p*, on the *i*-th observation of stimulus level, *l*. Since the time interval varied across cells, depending on how long we maintained stable recordings, incorporating this information into our model gave greater weight to data points with longer time intervals.

For levels less than threshold, firing rate was expected to be equal to the baseline firing rate. For stimulus levels greater than threshold, the cell could have an excitatory (*<u>s_c_</u>* > 0) or inhibitory (*s_c_* < 0) response.

The distribution of baseline rates across the cells in the small pupil condition were modeled by a Gamma prior, *b_c_*_,0_ ∼ *Gamma*(*b_α_*, *b_β_*), with *b_α_* ∼ *Gamma*(0.5,0.1) and *b_β_*∼ *Gamma*(0.1,0.1)^1^. We set the priors for *α* and *β* after inspecting the distributions of spontaneous rates in the small and large pupil condition. Although prior work (McGinley et al., 2015a) suggests that we would expect to see a difference in spontaneous rate between the pupil conditions, we chose an unbiased prior for the ratio (*b_c,1_*⁄*b_c_*_,0_) such that the mean and standard deviation were *Normal*(1, 1) and *HalfNormal*(1), respectively.

The distribution of slopes across cells were modeled using a Normal prior, *s_c_*_,0_ ∼ *Normal*(*s_μ_*, *s_σ_*), and the priors for the population mean and standard deviation were modeled themselves as *s_μ_*∼*Normal*(0.1, 0.1) and *s_σ_*∼ *HalfNormal*(0.1). The distribution of thresholds across cells was modeled using a Normal prior, *θ_c_*_,0_ ∼ *Normal*(*θ_μ_*, *θ_σ_*) with priors for the population mean and standard deviation modeled as *θ_μ_*∼ *Normal*(40, 5) and *θ_σ_*∼ *HalfNormal*(5). To minimize potential bias, unbiased priors were used for parameters representing the difference between pupil conditions. Specifically, the difference in slope (*s_c_*_,1_ − *s_c_*_,0_) was modeled with a Normal prior with mean and standard deviation of *Normaal*(0, 0.1) and *HalfNormal*(0.1), respectively. The difference in threshold (*θ_c_*_,1_ − *θ_c_*_,0_) was modeled with a Normal prior with mean and standard deviation of *Normal*(0, 5) and *HalfNormal*(5), respectively.

### Data analysis: Frequency-tuning curves

To quantify changes in frequency tuning linked to internal state, we first found best level (i.e., the level that evoked the maximum rate across all frequencies) using the FRA measured across all pupil-size bins. We then extracted frequency tuning curves (FTCs) at best level from the large-pupil and small-pupil FRAs. FTCs were smoothed by averaging across best level and the two neighboring levels in the FRA.

We assumed that the FTC for individual cells, *c*, and pupil condition, *p*, could be expressed as a Gaussian function with four parameters, baseline firing rate, *b*, gain, *g*, best frequency, *BF*, and band-width, *BW*:

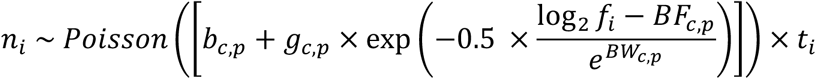

Here, *n* is the evoked rate for the cell, pupil condition (small vs. large) and stimulus frequency, *f*, on the *i*-th observation. As in the rate-level model, the cell’s firing rate during the 50 ms preceding sound on-set was a better measure of the cell’s baseline response for sub-threshold levels and the spontaneous rate model was also incorporated into this model to improve our estimate of *b*.

The distribution of gains across cells was modeled using a Normal prior, with priors for the population mean and standard deviation modeled as *g_μ_*∼ *Normal*(10, 1) and *g_σ_* = *HalfNormal*(10). The distribution of best frequency across cells was modeled using a Normal prior, with priors for the population mean and standard deviation modeled as *BF_μ_*∼ *Normal*(10, 1) and *BF_σ_* = *HalfNormal*(1). The distribution of bandwidth across cells was modeled using a Normal prior, with priors for the population mean and standard deviation modeled as *BW_μ_* ∼ *Normal*(−0.5, 0.1) and *BW_σ_* = *HalfNormal*(0.25).

To avoid bias in estimates of pupil-related changes, uninformative Normal priors were used for parameters representing the difference between small and large pupil conditions. Specifically, the logarithm of the ratio in gain (*g_c_*_,1_⁄*g_c_*_,0_) was a with a mean and standard deviation of *Normal*(0, 0.1) and *HalfNormal*(0.1), respectively. The logarithm of the ratio in bandwidth (*BW_c_*_,1_⁄*BW_c_*_,0_) had a mean and standard deviation of *Normal*(0, 0.1) and *HalfNormal*(0.25), respectively. For the ratio in gain and ratio in bandwidth, exp(0) = 1, such that the expected value of the prior was no effect of pupil size on gain or bandwidth. The difference in best frequency (*BF_c_*_,1_ − *BF_c_*_,0_) had a mean and standard deviation of *Normal*(0, 0.1) and *HalfNormal*(0.1), respectively.

### Data analysis: Sleep states

To test for neural correlates of sleep states, we defined a binary variable that indicated the presence or absence of sleep (sleep_i_(*t*, *s*)) and added it as a term in the baseline-gain model (see above),

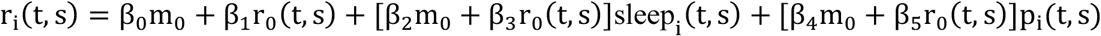

We also tested a sleep-only model in which baseline and gain were modulated only by sleep state,

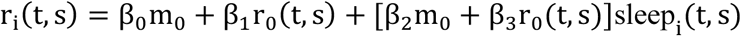

Model fitting and evaluation were completed as for the baseline-gain model.

### Data analysis: Local-field potential

The local-field potential (LFP) signals from each recording were bandpass filtered using a forward and reverse filtered second-order Butterworth filter into delta (1-4 Hz), theta (4-7 Hz), alpha (7-14 Hz), beta (15-30 Hz), low gamma (30-60 Hz) and high gamma (60-100 Hz) bands (Yuzgec et al., 2018). The instantaneous amplitude of each bandpass-filtered signal was calculated by computing the magnitude of its Hilbert transform, low-pass filtering with a cut-off at 1 Hz, and taking the mean amplitude across all electrodes.

### Experimental design and statistical analyses

Statistical tests used to assess whether model fits were significantly better than chance (i.e., whether they accounted for any auditory response, tuning parameter, or state-related changes in neural activity), and to compare predictions across models are described above, together with other aspects of the models. Tests for correlations between data across conditions are reported as the unsquared Pearson correlation coefficient (*r*) and a *t*-test for significance. Tests for differences between conditions are reported using two-tailed Wilcoxon rank-sum tests (*rank-sum test*), Wilcoxon signed-rank tests (*sign test*), or *t*-tests (*paired t-test* or *unpaired t-test*).

### Code

Custom software for stimulus presentation and pupil analysis is available at https://bitbucket.org/lbhb/baphy. MATLAB and Python code for statistical analyses is available from the corresponding author on request.

## Results

### Dilated pupil is correlated with increases in the spontaneous activity, gain, and reliability of auditory neurons

To examine the relationship between internal brain state and neural representation of natural sounds, we simultaneously recorded infrared (IR) video of the eye and single-unit spiking activity in the primary auditory cortex (A1) of head-restrained, non-anesthetized ferrets (Fig 1a, *n* = 114 neurons from 46 recording sites in 3 ferrets). Pupil size could be extracted from the video, providing a measure of internal state (Reimer et al., 2014; McGinley et al., 2015a; Vinck et al., 2015). In order to gather data on the response to the same sounds across a wide range of states, our stimulus consisted of a small number of sounds (two ferret vocalizations) repeated many times (up to 120 repetitions per recording, mean +/− SD = 77 +/− 40 repetitions). To further increase our sampling of internal states, we paused this presentation at set times in the recording to rouse the ferret using auditory stimuli (see Materials and Methods). The variability of neural activity across repetitions of the same vocalization segments exceeded what would be predicted by models of spiking as a Poisson process (Fig. 1b-d, mean Fano factor +/− SEM = 1.5 +/− 0.07). This degree of variability is consistent with previous in vivo studies in cortex (Shadlen and Newsome, 1998; Churchland et al., 2010; Goris et al., 2014).

**Fig. 1:**
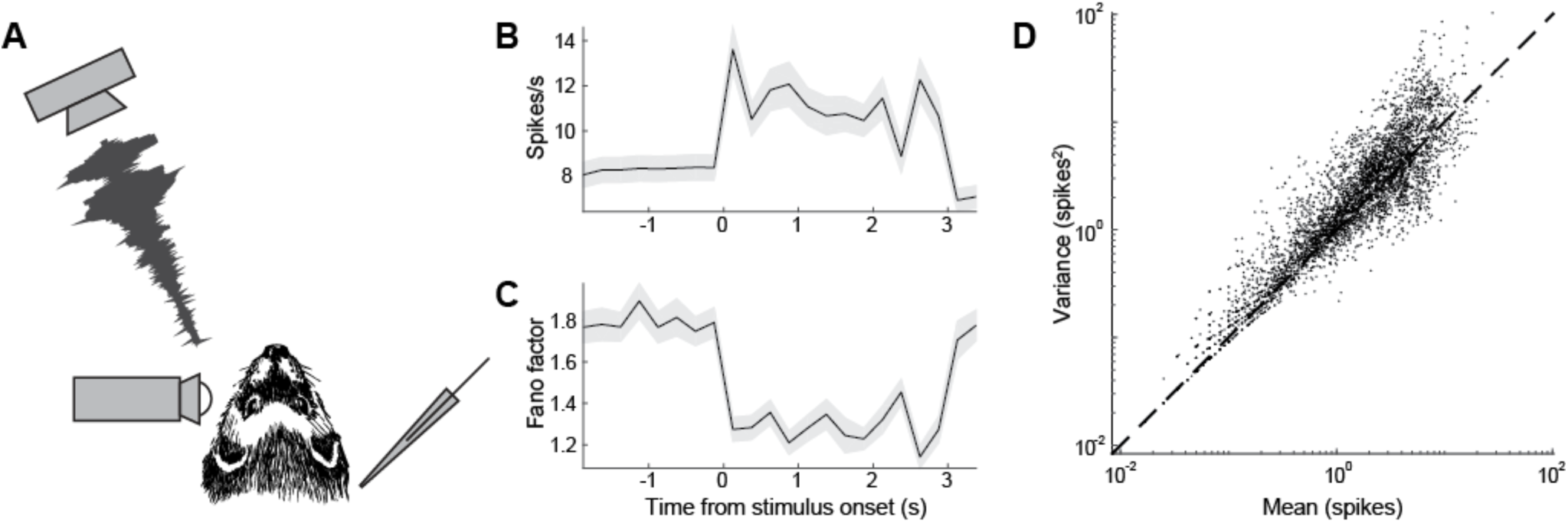
Ferret auditory cortex shows trial-to-trial variability in neural responses to sound. (A) Schematic of experiment, illustrating ferret, free-field speaker, camera recording video of eye, and contralateral extracellular electrode. (B) Mean response to sound across all neurons in the vocalization dataset (*n* = 114). Shading indicates SEM. (C) Time-varying Fano factor for all neurons in the vocalization dataset. Shading indicates SEM. (D) Variance-to-mean relationship for all neurons in the vocalization dataset. Each data point illustrates the mean and variance of the spike count of one neuron during one 250-ms segment of one vocalization or the surrounding intervals. Dashed line indicates prediction for a Poisson process (slope = 1).

Although lighting conditions were held static during recordings, we also observed large changes in pupil size across timescales of tens of seconds to minutes (Fig. 2a). Visual inspection of spike rasters suggested that trial-to-trial variability in neural activity sometimes tracked changes in pupil size (Fig. 2b). To quantify the association between pupil size and firing rate, we fit a linear model that allowed baseline spike rate and gain of sound-evoked responses to depend on pupil size (baseline-gain model, see Materials and Methods, Fig. 2c). In a majority of neurons (*n =* 75/114, 66%) this model was significantly more accurate at predicting single-trial neural activity than a control model where pupil was shuffled in time (i.e., the prediction was based only on the neuron’s peristimulus time histogram [PSTH] response to each vocalization) (Fig. 2d, *p* < 0.05, permutation test). The degree of improvement was not correlated with the prediction accuracy of the control model alone (*r* = −0.08, *p* = 0.5, *t*-test), suggesting that the influence of pupil on trial-to-trial variability did not depend on a neuron’s overall auditory responsiveness.

**Fig. 2:**
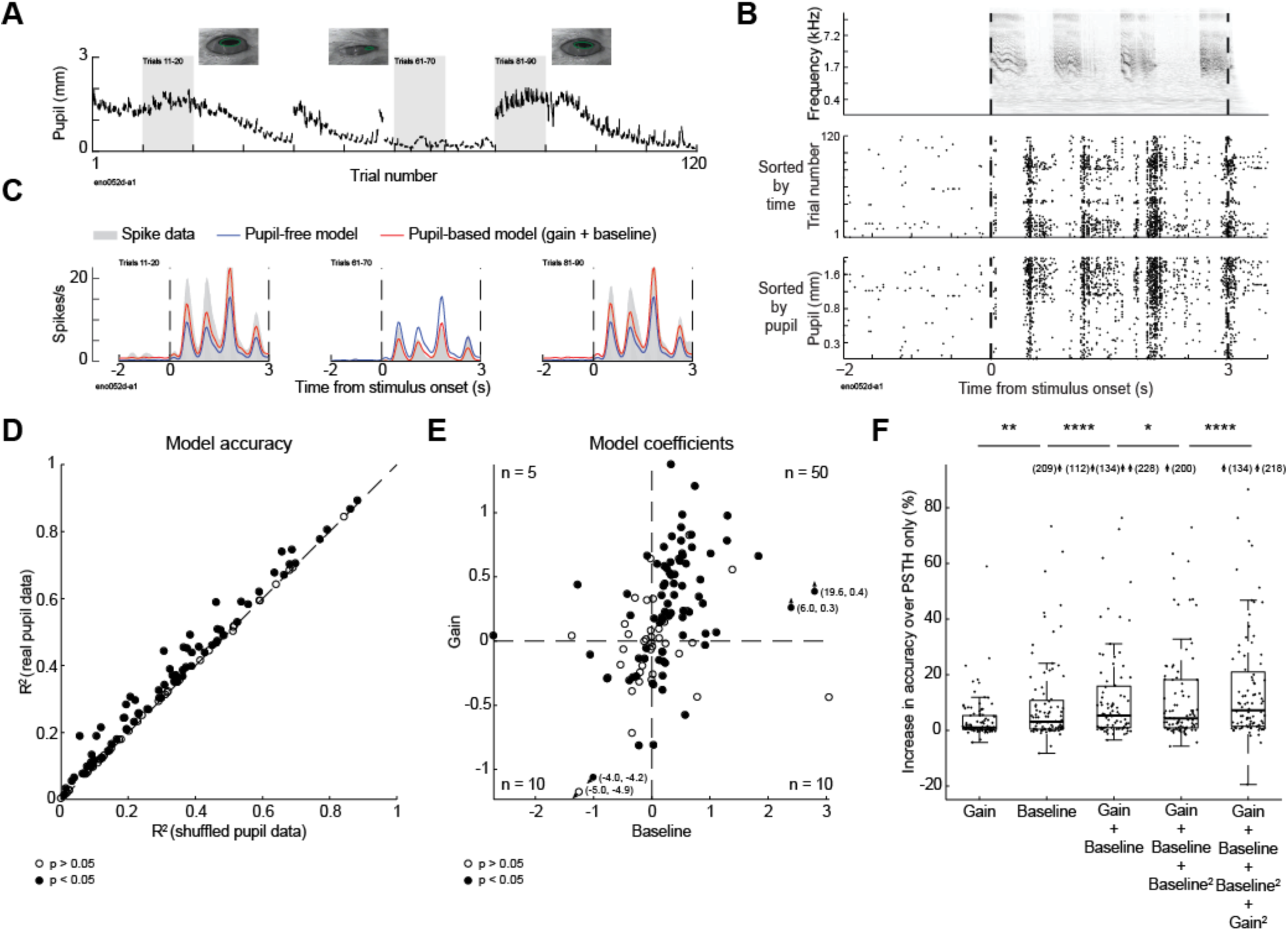
Trial-to-trial variability in neural responses to sound tracks changes in pupil size. (A) Pupil trace recorded across 120 repetitions of one ferret vocalization, with example video frames. (B) Spectrogram of one ferret vocalization and spike raster of one neuron’s response to the vocalization, sorted by time (top) and prestimulus pupil size (bottom). (C) Predicted and actual pupil-dependent changes in spiking activity for the neuron shown in 2a-b. Panels show peristimulus time histogram (PSTH) responses averaged across blocks of 10 stimulus presentations selected from epochs with different pupil size (see 2a). PSTHs predicted by the null model (blue) and baseline + gain model (red) are overlaid on the actual PSTH (gray shading). (D) Accuracy of pupil-based baseline-gain regression model for neural responses to ferret vocalizations (*n* = 114 neurons), plotted against accuracy of a control, PSTH-only model, fit to temporally shuffled pupil data from each cell. Filled dots indicate neurons with a significant improvement over the control model (*n* = 75/114, *p* < 0.05, permutation test). (E) Coefficients of baseline-gain model for all data from vocalization recordings (*n* = 114 neurons). Positive values indicate an increase in baseline spike rate (horizontal axis) or response gain (vertical axis) with larger pupil. Numbers in corners of each quadrant indicate the count of neurons with parameters in that quadrant and a significant improvement in prediction accuracy over the control, pupil-independent model. (F) Improvement in accuracy over PSTH-only model for various models for population of neurons with a significant fit for any model (*n* = 92/114, *: *p* < 0.05, **: *p* < 0.01, ****: *p* < 0.0001, sign test). Boxplot indicates median, interquartile range, and 1.5 times the interquartile range.

The baseline-gain model included separate terms for effects of internal state on gain and baseline firing rate, both of which tended to be positive, indicating an increase in firing rate with increasing pupil size (Fig. 2e). Both the baseline and gain terms contributed to model prediction accuracy, compared to models that included only one term (Fig. 2f, Table 1). Adding second-order terms improved median prediction accuracy further (Table 1), and increased the number of neurons that showed a significant fit from 75 to 79.

**Table 1.**
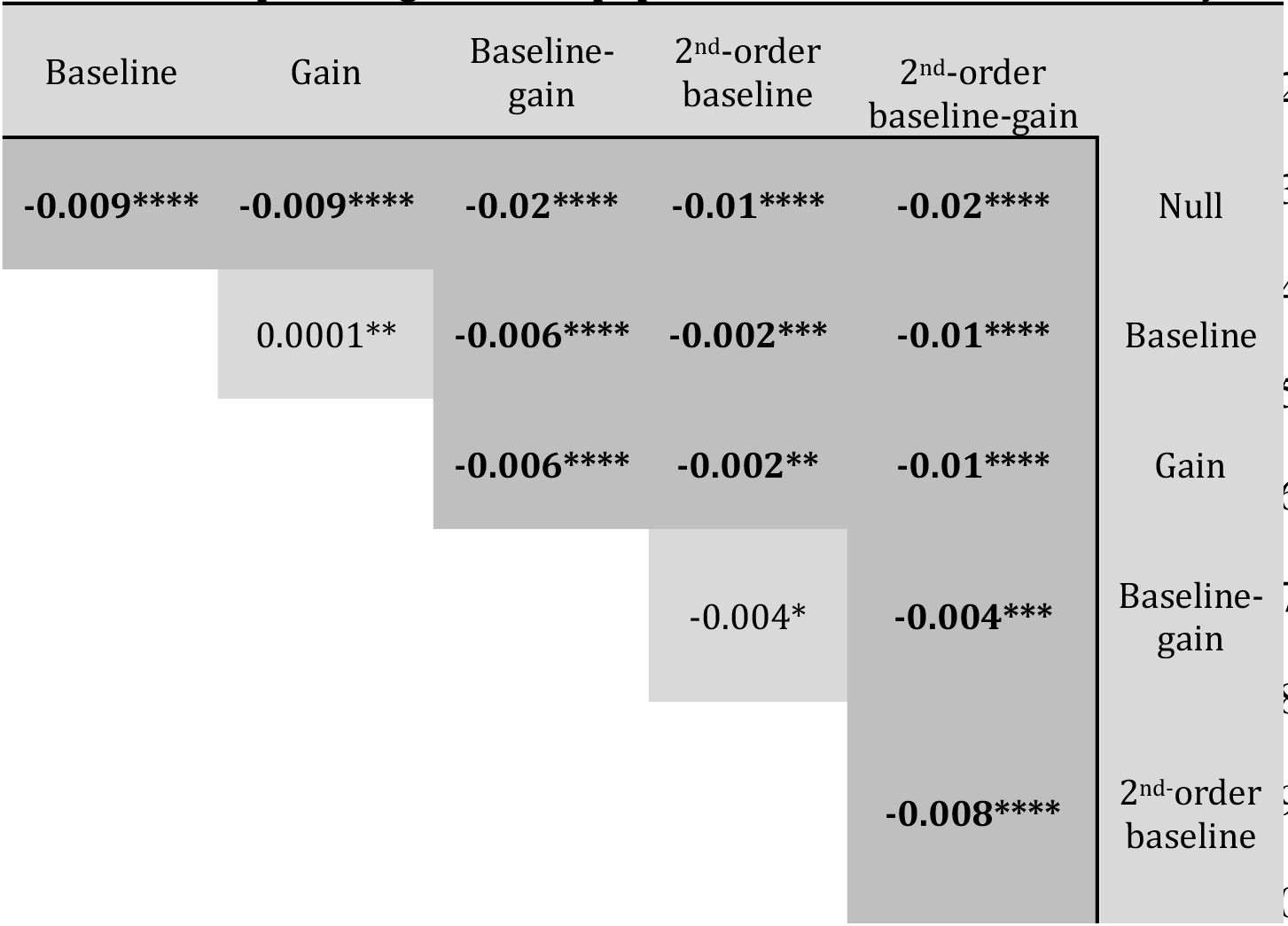
Stepwise regression of pupil-based models of neural activity. Stepwise comparison of accuracy of different pupil-based models of neural activity applied to vocalization dataset. Numbers indicate difference in median accuracy across the population of neurons with significant fit for any model (*n* = 92/114). Bold text indicates comparisons for which the difference is significant at alpha level 0.05 with Bonferroni correction for multiple comparisons (*: *p* < 0.05, **: *p* < 0.01, ***: *p* < 0.001, ****: *p* < 0.0001, sign test).

Given that pupil size was more variable in some recordings than others, we wondered whether we were unable to detect effects of internal state in some neurons simply because the animal’s pupil size remained fairly stable during the recording. Variability of pupil size during the recording was greater for neurons that showed a significant effect of pupil (median SD = 0.4 mm, *n* = 79 neurons) than for neurons that did not (median SD = 0.2 mm, *n* = 35 neurons, *p* = 1e-5, rank-sum test). Among neurons that showed a significant effect of pupil on firing rate, there was a correlation between the variance of pupil during the recording and the degree of improvement of the model over a model based on PSTH response alone (*n* = 79 neurons, *r* = 0.37, *p* = 4e-4, *t*-test). Thus, the measurement of 66% of neurons showing a significant effect of state on activity represents a lower bound on the frequency of neurons showing pupil-related effects. However, splitting the data at the median pupil size observed during each recording revealed a consistent shift towards high-frequency local-field potential (LFP) activity when pupil was large (Fig. 3), echoing previous work in mouse visual cortex and suggesting that median pupil size could be used to distinguish high and low-arousal states (Harris and Thiele, 2011; Vinck et al., 2015).

**Fig. 3:**
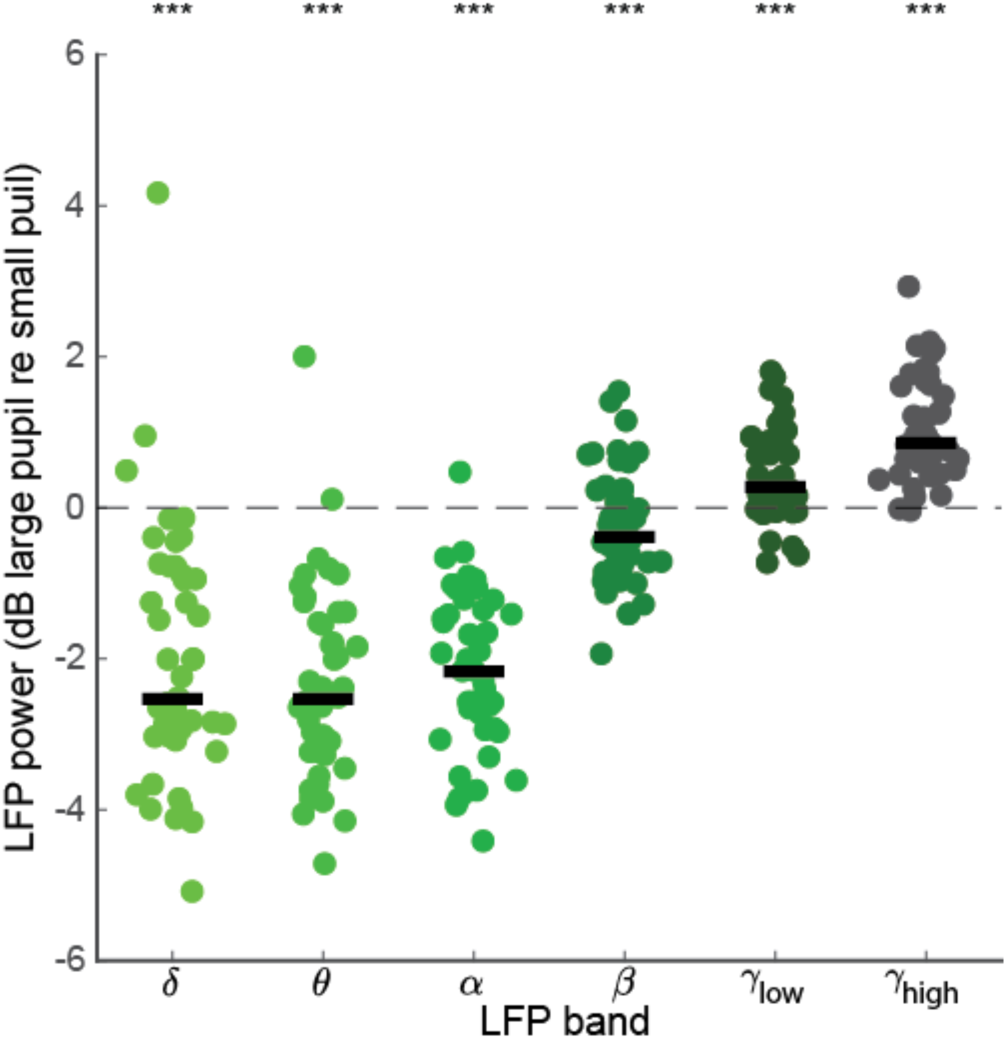
Dilated pupil is associated with neural desynchronization. Log ratio change (dB) in local field potential (LFP) power between recording segments with large and small pupil (*n* = 46 recordings). Thick line indicates population median (***: *p* < 0.001, rank-sum test on hypothesis that median is equal to 0). Putative sleep states (see Fig. 8) have been excluded from the comparison.

The models were fit to data that included sound-evoked activity as well as activity during the silent intervals before and after each vocalization. The baseline term of the models could reflect either modulation of spontaneous activity or a stimulus-dependent modulation of sound-evoked firing rate that did not scale with stimulus strength. To distinguish between these possibilities, we fit a model with only a baseline term to data from the prestimulus epoch alone, ignoring sound-evoked activity. We then fit a model that included terms for both baseline and gain to the peristimulus epoch alone, ignoring spontaneous activity. The magnitude of the baseline terms in the two models was correlated (*n* = 75 neurons, *r* = 0.55, *p* = 4e-08, *t*-test), suggesting that the baseline term of the model reflected primarily modulation of spontaneous activity rather than (or in addition to) stimulus-evoked activity.

Previous research indicates that the level of pupil-indexed arousal is associated with changes in the reliability of neurons in auditory cortex, where reliability is defined as the mean cross correlation between spiking responses to the same stimulus across repetitions (Reimer et al., 2014; McGinley et al., 2015a). To test for this effect, we calculated the correlation between the spiking response evoked by each repetition of each vocalization (Fig. 4a). Inter-trial correlations increased with prestimulus pupil size, indicating a more reliable response to sound (Fig 4a-b). To quantify the change in reliability, we split data into large-pupil and small-pupil trials based on whether mean pupil diameter in the silent interval preceding each vocalization was greater or less than the median pupil size during the recording (Fig. 4c). Across the entire population, reliability was significantly greater during large-pupil trials (Fig. 4c, *n* = 114 neurons, *p* = 5e-8, paired *t*-test). This difference in reliability was preserved in the subset of cells that showed modulation of neural activity by pupil-indexed state under our second-order baseline-gain model (Fig. 4d, *n* = 79 neurons, *p* = 5e-8, paired *t*-test), but disappeared when comparing only cells that did not show pupil-associated modulation under this model (Fig. 4d, *n* = 35 neurons, *p* = 0.2, paired *t*-test). These results suggest that pupil-associated state affects both the mean rate at which auditory neurons respond to sound and the variability of their responses, and that these effects occur in the same population of cells.

**Fig. 4:**
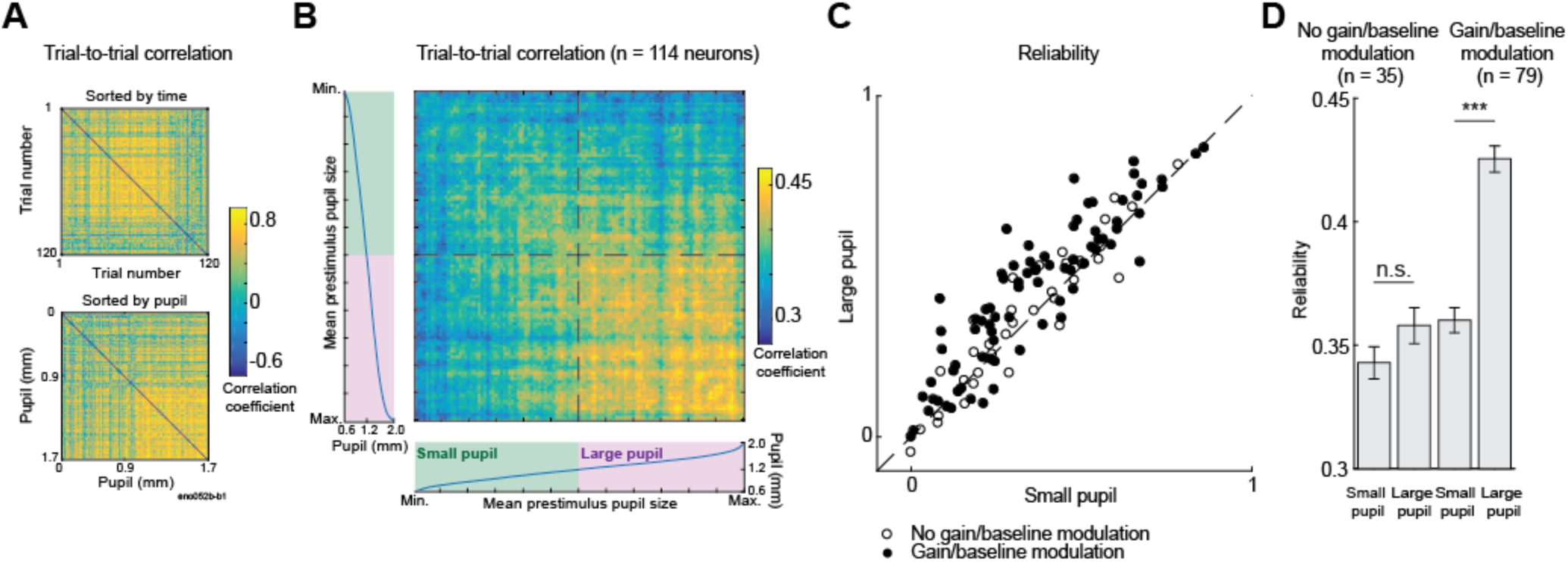
Reliability of neural response to sound increases when pupil is dilated. (A) Correlation between time-varying activity across trials evoked by one ferret vocalization in one neuron, sorted by trial order (top) and mean pupil size before stimulus onset (bottom). (B) Mean trial-to-trial correlation for all neural responses to ferret vocalizations. This heat map was constructed by computing the average correlation matrix sorted by pupil size across all neurons (*n* = 114). For display, the correlation of each trial with itself was replaced by the mean correlation for the two trials with the most similar pupil size before averaging. Marginal plots indicate the mean pupil size in each trial, across all neurons. Dashed line indicates median pupil size. (C) Comparison of reliability (mean trial-to-trial correlation) for each neuron’s response to vocalizations (*n* = 114) for small versus large pupil. Trials are classified as “large pupil” and “small pupil” based on whether the mean pupil size preceding the vocalization is greater or less than the median pupil size during the recording. Filled circles indicate cells with a significant fit under a second-order baseline-gain regression model (*n* = 79, *p* < 0.05, permutation test). (D) Mean reliability (+/− SEM) in each condition for subpopulations of cells that do or do not show effect of pupil-associated state under the regression model (***: *p* < 0.001, paired t-test).

### Non-monotonic effects of pupil on firing rate are observed in some A1 neurons

Arousal can have a non-monotonic effect on behavior: both learning (Yerkes and Dodson, 1908; Diamond et al., 2007) and task performance (Aston-Jones and Cohen, 2005; Murphy et al., 2011; van Kempen et al., 2019) are sub-optimal in minimally and maximally aroused states. A previous study found that multiunit spiking activity in mouse auditory cortex showed similarly non-monotonic relation between pupil and firing rate: on average, spontaneous activity was lowest, and gain highest, at intermediate pupil diameter, suggesting that this range of pupil sizes was optimal for detection of auditory signals (McGinley et al., 2015a). We found examples of non-monotonic pupil-firing rate relationships in some cells (Fig. 5a). To test for their prevalence, we fit data on neural responses to ferret vocalizations with a segmented regression model (Simonsohn, Uri, 2018) that predicted a linear effect of pupil on neural activity but allowed the slope of the fit to take two different values depending on pupil size (Fig. 5b). A difference in the sign of the slope between segments indicated a non-monotonic relationship between pupil size and spiking activity. We then compared the accuracy of this model to a similar model in which slope could vary between segments, but both line segments were constrained to have the same sign (i.e., the fit could not assume the shape of a U or inverted U).

**Fig. 5:**
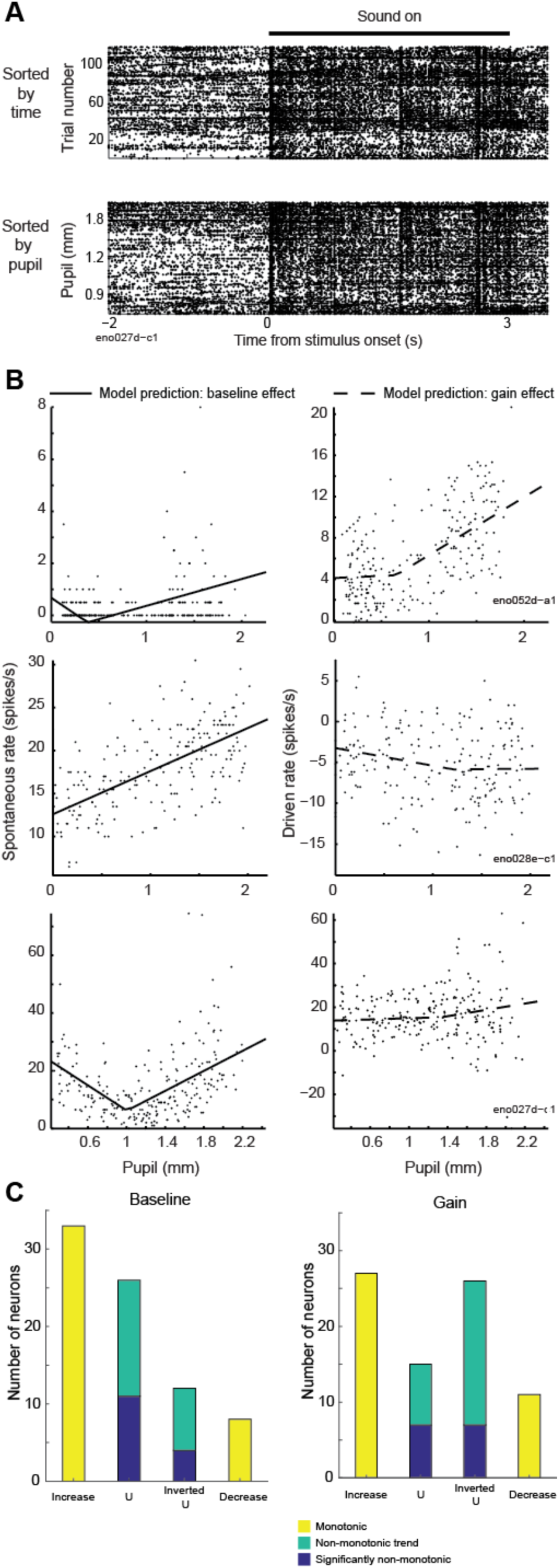
Non-monotonic effects of pupil-related state. (A) Spike raster of one neuron’s response to single ferret vocalization sorted by time (top) and pupil size preceding the stimulus (bottom). A non-monotonic relationship is evident between pupil size and spontaneous firing rate. (B) Examples of segmented regression model fits to spontaneous and driven activity for three neurons. Each point represents spike rate for one neuron on one trial (spontaneous activity: 2 s, driven activity: 3 s). The bottom row displays data from the neuron in panel A. (C) Results of test for non-monotonicity using segmented regression model, counting the number of neurons with significant or trends toward non-monotonic changes in baseline or gain.

Although a large proportion of neurons that showed effects of pupil-associated state also showed a trend towards non-monotonicity in either baseline firing rate or gain (Fig. 5c, *n* = 38/79 [baseline], *n* = 41/79 [gain]), allowing for a non-monotonic segmented fit significantly improved the accuracy of the model for a smaller fraction of cells (*n* = 15/79 [baseline], 14/79 [gain], *p* < 0.05, permutation test). Among neurons that showed a trend towards non-monotonicity, more showed trends towards U in baseline firing rate than inverted U, and more showed an inverted U than U in gain (Fig. 5c), suggestive of previous results comparing spontaneous and sound-evoked activity in mouse A1 (McGinley et al., 2015a).

### Frequency and level tuning show no or small dependence on pupil size

To examine the effect of changes in internal state on stimulus selectivity, we recorded the responses of A1 neurons to tone pips at a variety of frequencies and levels (Fig. 6a-c, 114 neurons from 40 recording sites in 4 ferrets). To test for changes in frequency and level selectivity, we divided the data from each cell into large-pupil and small-pupil bins based on the median pupil size observed during the recording (Fig. 6a-b), and constructed frequency response areas (FRAs) from the data in each bin (Fig. 6c). FRAs showed a variety of patterns typical of the auditory system (Bizley et al., 2005), including a sound-level response threshold and broadening of spectral bandwidth at higher sound levels (Fig. 6c).

**Fig. 6:**
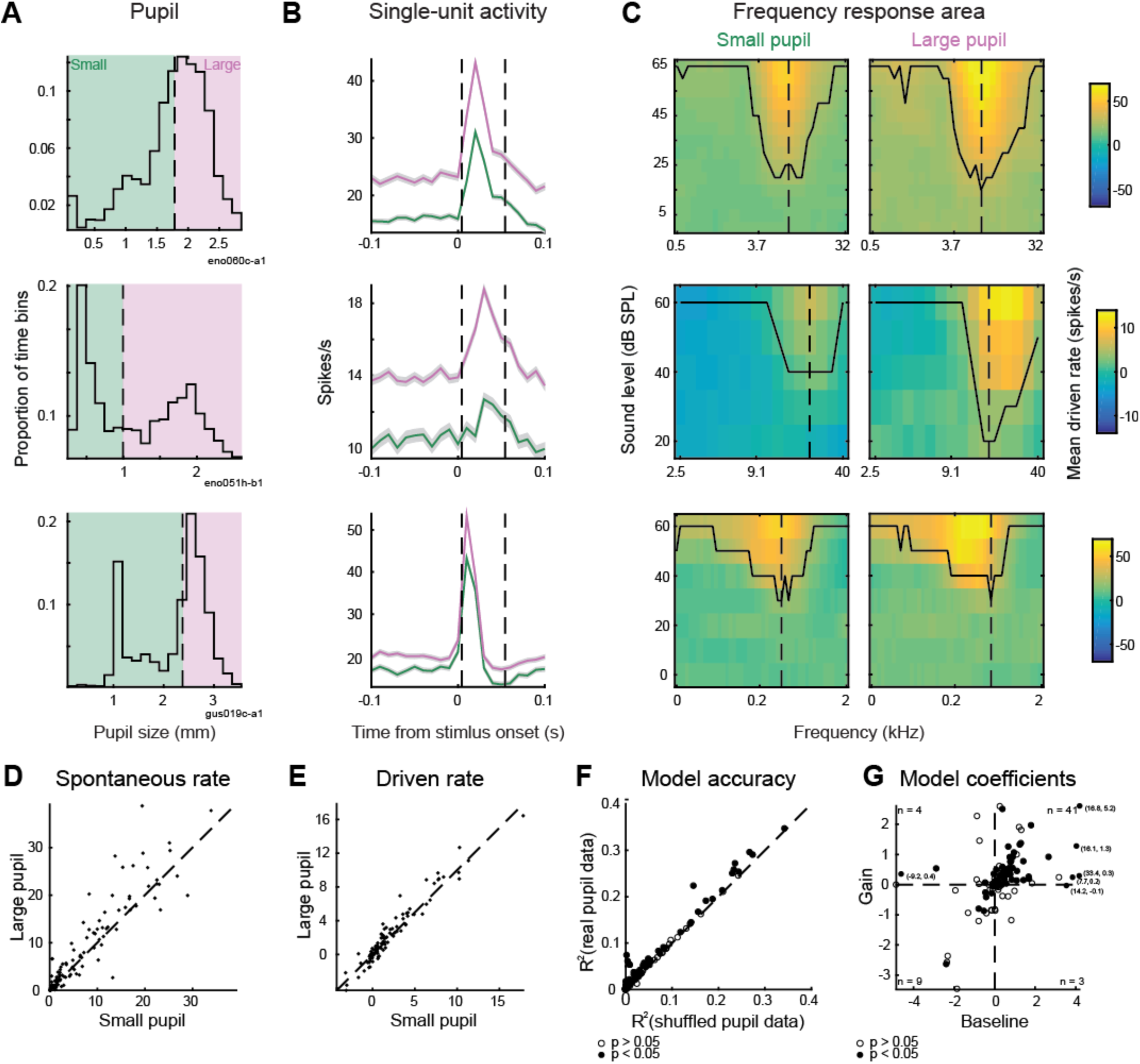
Spontaneous activity and evoked response to tones increases when pupil is dilated. (A) Distribution of pupil size during three recordings of neural responses to tone pips, indicating division into small-pupil and large-pupil bins based on median pupil size during the recording. (B) Mean peristimulus time histograms (PSTH)s of neural responses for the three example cells, computed separately for data in the smal-pupil (green) and large-pupil (purple) bins and averaged across all tone pip levels and frequencies. Shading indicates SEM. Dashed lines indicate window for measurement of response to sound. (C) Frequency-response areas for the three examples cells, computed separately for small and large pupil bins. Contour indicates lowest level that shows an auditory response at each frequency. Dashed line indicates characteristic frequency. (D-E) Spontaneous and driven rate for all neurons’ responses to tone pips (*n* = 114 neurons). (F) Single-trial prediction accuracy of linear baseline-gain model and control, pupil-independent model for all neurons’ response to tone pips. Filled dots indicate neurons with a significant improvement for the baseline-gain model (*n* = 57/114, *p* < 0.05, permutation test). (G) Coefficients of baseline-gain model fit for all data from tone-pip recordings. Numbers indicate count of neurons with significant improvement for the pupil-dependent model in each quadrant.

Comparing data from the large and small-pupil bins showed a systematic increase in spontaneous activity and the response to tones across all levels and frequencies (Fig. 6b,d-e, *n* = 114 neurons*, p* = 1e-4 [spontaneous rate], *p =* 0.03 [driven rate], sign test). To confirm that this result did not depend on our division of the data into large and small-pupil bins, and to further examine changes in driven rate, we used linear regression to predict the response to tones based on pupil size and the average FRA of the neuron (see Materials and Methods). As it did for vocalizations, pupil size predicted trial-to-trial variability in a subpopulation of cells (Fig. 6f, *n* = 57/114 neurons, 50%, *p* < 0.05, permutation test). Most state-modulated neurons showed enhanced gain and baseline firing rate when pupil was large (Fig. 6g), suggesting that pupil-indexed state acted to shift and multiplicatively scale the neuron’s auditory tuning curve.

Does the stimulus selectivity of auditory neurons change depending on pupil-indexed state? To test for changes in level tuning, we again divided data at the median pupil value in each recording, then fit a hinge function (see Methods and Materials) to the characteristic-frequency rate-level function in each pupil condition (Fig. 7a-b). Across the population, there was a significant decrease in threshold when pupil was large (mean change: −2.65 dB, 90% credible interval: −4.00 to −1.33 dB), but no change in the slope of the rate-level function (mean change: 0.05 [spikes/s]/dB, 90% credible interval: −0.07 to 0.15 [spikes/s]/dB SPL). Consistent with previous analyses, we also observed an increase in the baseline firing rate of the cells when pupil was large (mean change: 1.68 spikes/s, 90% credible interval: 0.80 to 2.59 spikes/s). At the single-cell level, most neurons showed a significant change in baseline firing rate (*n* = 68/114), but few showed significant changes in slope (*n* = 12/114) or threshold (*n* = 8/114). The low number of cells showing significant changes in threshold may be due to the small effect size (−2.65 dB average across the population) and the uncertainty in model fits for individual cells.

**Fig. 7:**
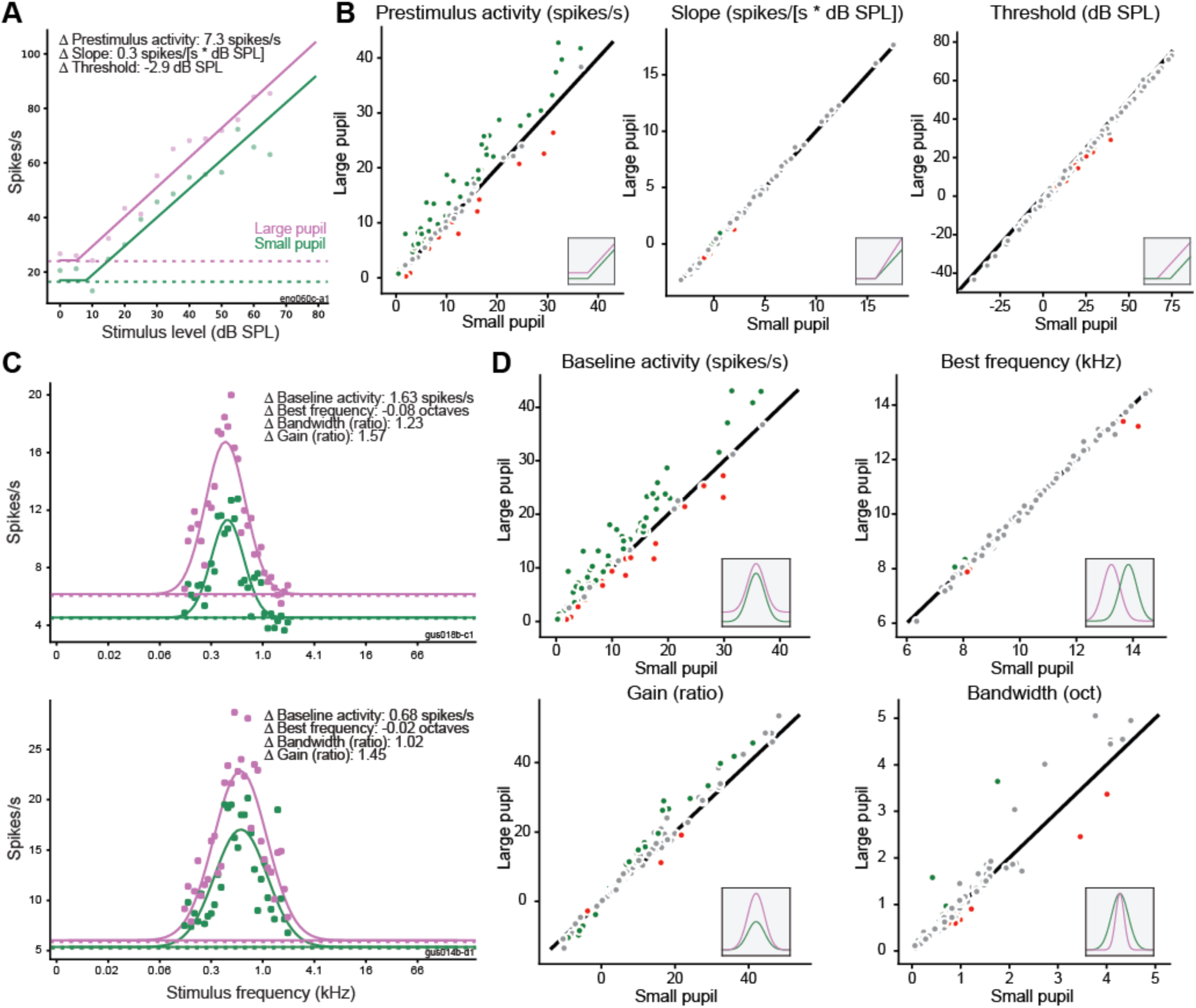
Effects of pupil-related state on frequency and level tuning. (A) Example of hinge function fit to neural response at characteristic frequency for large-pupil (purple) and small-pupil (green) conditions. Dashed lines indicate prestimulus firing rate in each condition. (B) Change in each parameter of rate-level function between pupil conditions (large pupil vs. small pupil) for all neurons (*n* = 114). Inset cartoons illustrate effect of changing each parameter fit. Green and red dots indicate neurons with a significant change in the tuning parameter (90% credible interval for difference between conditions outside 0). Gray dots indicate neurons with changes that were not significant at the single-cell level. (C) Examples of Gaussian functions fit to neural responses at best level for large-pupil (purple) and small-pupil conditions (green). (D) Change in each parameter of frequency-tuning curve across pupil conditions for all neurons (*n* = 114). Insets and color coding as in panel B.

To test for changes in frequency tuning, we fit a Gaussian function to the best-level frequency tuning curve (FTC) in each pupil condition (Fig. 7c-d). To isolate changes in frequency tuning (i.e., bandwidth) from nonspecific changes in baseline firing rate or gain, our model included multiplicative gain term and additive offset terms. Consistent with data from previous analysis, the gain and baseline parameters of the FTC showed a systematic increase across the population when pupil was large (gain: mean large/small ratio = 1.12, 90% credible interval = 1.06 to 1.18, baseline: mean change = 2.02 spikes/s, 90% credible interval = 1.20 to 2.75). There was no change in the spectral bandwidth of the FTC when pupil was large (mean large/small ratio = 1.01, 90% credible interval = 0.94 to 1.07). There was also no change in mean best frequency across pupil conditions (mean change = −0.02 octaves, 90% credible interval = −0.09 to 0.05). The number of neurons that showed a significant change in each parameter was greater for gain (*n* = 27/114) and baseline firing rate (*n* = 78/114) than it was for bandwidth (*n* = 14/114) or best frequency (*n* = 4/114). Thus, while A1 neurons did sometimes show pupil-dependent changes in response threshold or other tuning parameters, these changes were relatively small compared to the changes in baseline activity and response gain.

### Sleep states account for additional neural variability

Although we did not explicitly seek to study sleep state, during some recordings we observed periods of tonically constricted pupil accompanied by an increase in saccade-like eye movements (Fig. 8a-b, Movies 1 and 2). We initially speculated that these bouts represented rapid eye-movement (REM) sleep, based on a previous report correlating constricted pupil and REM sleep in mice (Yuzgec et al., 2018). However, the delta/theta ratio of local-field potential (LFP) – a signature of REM sleep (Yuzgec et al., 2018) – did not show a systematic change between putative REM bouts and recording segments with pupil size in the same range (*p* = 0.6, *n* = 26 recordings, rank-sum test). Instead, the increase in eye movements was accompanied by a decrease in alpha power (Fig. 8c, *p* = 1e-05, *n* = 26 recordings, rank-sum test), suggesting that the eye movements indicated sleep onset (Silber et al., 2007).

**Fig. 8:**
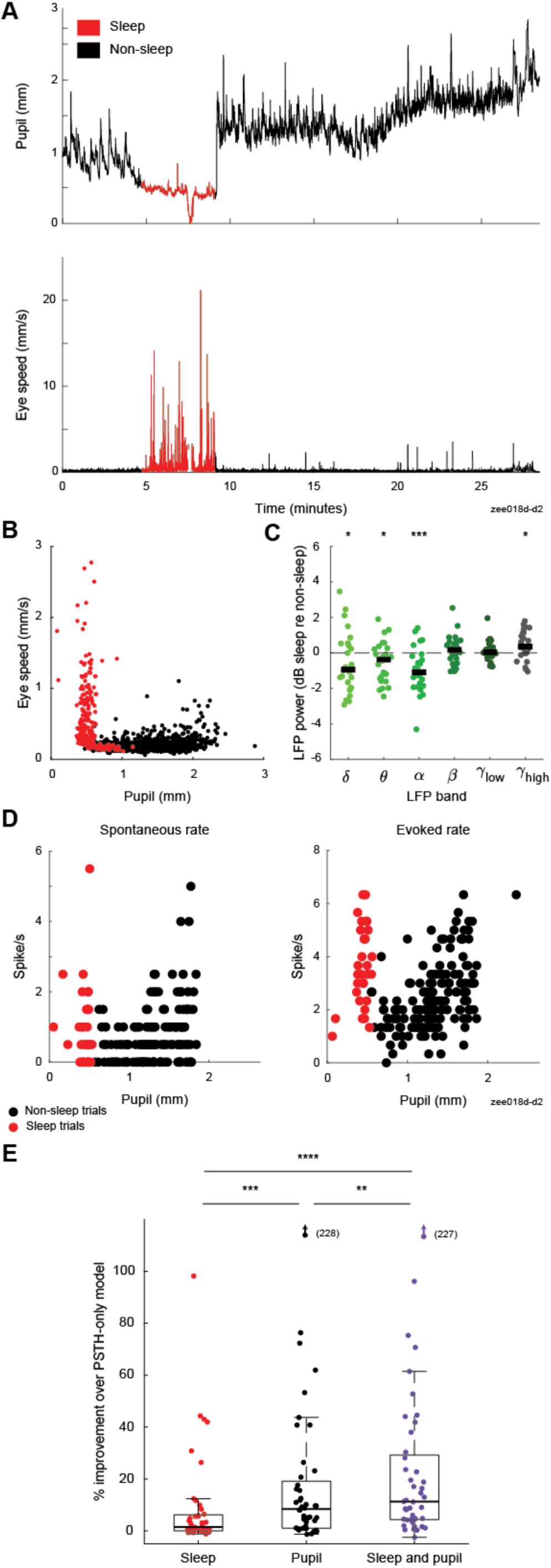
Sleep states account for additional variability in neural responses. (A) Pupil size and eye-speed traces from one recording. (B) Pupil size and eye speed data from one ferret (13 recordings). Each point illustrates average pupil size and eye speed during one trial (5.5 s). Color indicates whether a trial was classified as sleep (red) or non-sleep (black), using criteria based on eye movement and pupil size data (see Methods). (C) Relative change in LFP power between recording segments with and without sleep, matched for pupil size (*n* = 26 recordings). Bold line indicates population median (*: *p* < 0.05, ***: *p* < 0.001, rank-sum test on hypothesis that median is equal to 0). (D) Spontaneous (left) and sound-evoked neural data (right) from the recording shown in A. Each point indicates spike rate before or during one vocalization (prestimulus period = 2 s, stimulus period = 3 s). (E) Stepwise regression results for models including sleep state, pupil size or both sleep state and pupil size as predictors of spike rate (n = 44 neurons with significant fit for any model and recorded during experiments that include sleep episodes, **: *p* < 0.01, ***: *p* < 0.001, ****: *p* < 0.0001, sign test). Boxplot indicates median, interquartile range, and 1.5 times the interquartile range.

Given that neural responses to sound in auditory cortex are preserved during natural sleep states, but sometimes suppressed or enhanced (Issa and Wang, 2008), we wondered if these brief sleep episodes might show changes in firing rate distinct from those associated with changes in pupil size. We therefore fit linear regression models that included putative sleep state, pupil size, or both sleep state and pupil size as predictors of neural activity (Fig. 8c-d and Table 2). Sleep state significantly predicted neural activity in 81% of neurons recorded in experiments that included one or more sleep episodes (*n* = 38/47 neurons). Across the population of neurons with a significant fit for any tested model (*n* = 44), sleep state predicted less variability than pupil, which in turn predicted less variability than both pupil and sleep state (Fig. 8d and Table 2, median change in accuracy over PSTH-only model: 1.5% [sleep], 8.4% [pupil], 11% [pupil and sleep]). There was no correlation between the duration of sleep during the recording and the improvement in accuracy of the sleep-based model (*r* = 0.2, *p* = 0.16, *t*-test), suggesting that sleep effects vary across neurons.

**Table 2:**
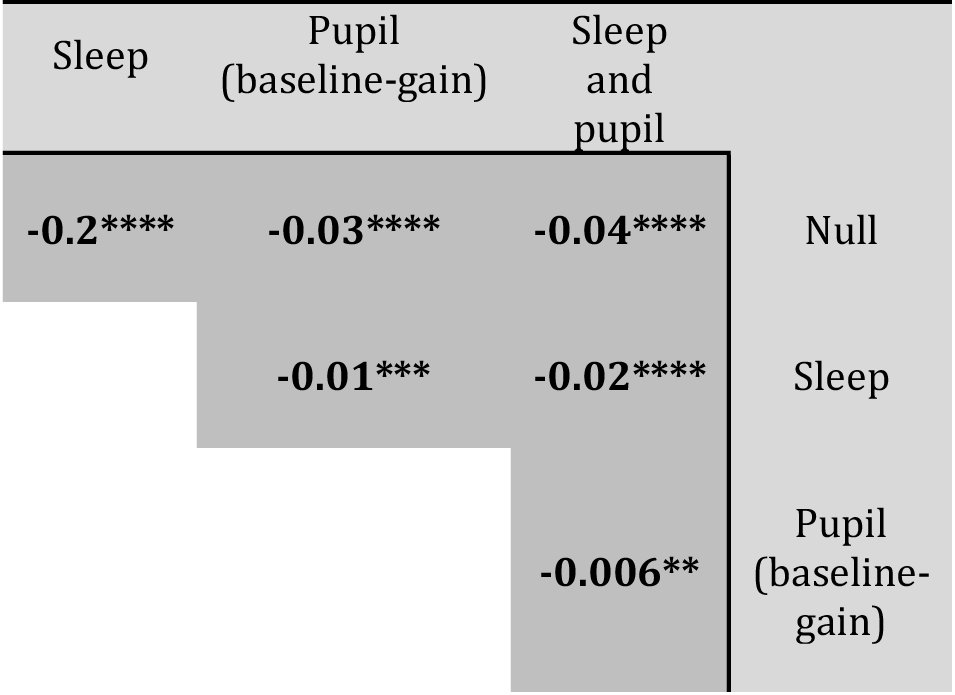
Stepwise comparison of pupil-and/or REM-based models of neural activity. Stepwise comparison of accuracy of pupil and/or sleep-state based models of neural activity (see Methods) applied to all vocalization data including sleep episodes (*n* = 47 neurons). Text indicates median accuracy for population of neurons with significant fit for any model (*n* = 44/47 neurons). Bold text indicates that the difference is significant at alpha level 0.05 with Bonferroni correction for multiple comparisons (*: *p* < 0.05, **: *p* < 0.01, ***: *p* < 0.001, ****: *p* < 0.0001, sign test).

| Sleep | Pupil<br>(baseline-gain) | Sleep<br>and<br>pupil |  |
| --- | --- | --- | --- |
| **-0.2\*\*\*\*** | **-0.03\*\*\*\*** | **-0.04\*\*\*\*** | Null |
|  | **-0.01\*\*\*** | **-0.02\*\*\*\*** | Sleep |
|  |  | **-0.006\*\*** | Pupil<br>(baseline-gain) |

In some neurons, visual inspection of spiking activity and pupil state indicated that sleep episodes were associated with a non-monotonic change in firing rates (Fig. 8c). To quantify this effect, we compared trials recorded during sleep episodes to non-sleep trials falling in an equivalent range of pupil sizes (Fig. 9a-b). Across the population, there was no significant difference in spontaneous rate, sound-evoked rate, or reliability between these two conditions (Fig. 9c, *p* = 0.16 [spontaneous rate], *p* = 0.38 [evoked rate], *p* = 0.17 [reliability], sign test, *n* = 47 neurons with recordings that included sleep episodes). However, a subpopulation of neurons did show a difference in spontaneous or sound-evoke rates at the single-cell level (Fig. 9c, *n* = 7/47 [spontaneous rate], *n* = 15/47 [evoked rate], *p* < 0.05, unpaired *t*-test). To further examine the effect of sleep state, we removed data from trials recorded during sleep episodes from the dataset and refit the segmented regression model initially used to test for non-monotonic effects. Fitting to data that excluded sleep state reduced the number of neurons that showed non-monotonic effects in both baseline and gain (Fig. 9d), indicating that sleep accounted for non-monotonic effects in some neurons (*n* = 7/42 [change in non-monotonicity of baseline effect], *n* = 6/42 [change in non-monotonicity of gain effect], *n =* 42 neurons with recordings that included sleep episodes and a significant fit for second-order baseline-gain model).

**Fig. 9:**
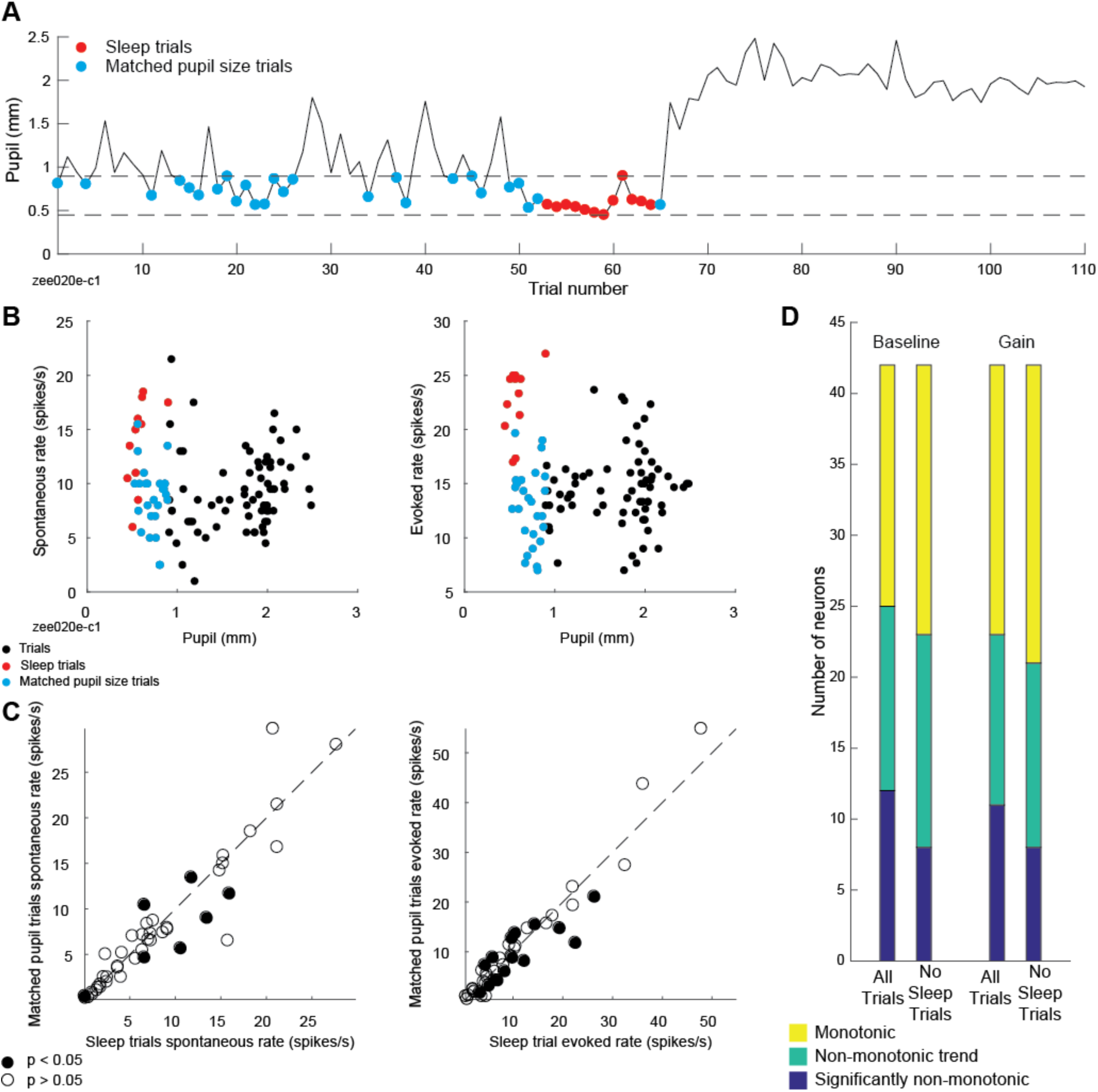
Sleep state accounts for non-monotonicity in some neurons. (A) Mean prestimulus pupil size for responses to one ferret vocalization, highlighting sleep trials (red) and trials with matched prestimulus pupil size (blue). (B) Spontaneous (i.e. prestimulus) and stimulus-evoked neural activity from the recording in A, illustrating greater spike rate in sleep trials as compared to trials with matched pupil size. (C) Average spontaneous (left) and sound-evoked spike rate (right) during sleep trials versus trials with matching pupil size (*n* = 47 neurons recorded in experiments including sleep episodes, filled circles: *p* < 0.05, unpaired t-test). (D) Results of test for non-monotonicity using segmented regression model fit to all data or data with sleep trials removed (*n* = 42 neurons recorded in experiments including sleep episodes, and showing significant fit to second-order baseline-gain regression model).

## Discussion

Pupil size is an indicator of central neuromodulatory processes related to arousal that affect processing in sensory cortex (McGinley et al., 2015b; Reimer et al., 2016). Our results show that pupil size is correlated with changes in the gain and baseline firing rate of neurons in the primary auditory cortex (A1) of non-anesthetized ferrets. Non-monotonic effects of changes in pupil-indexed state were observed in some neurons, but the majority of effects were monotonic and showed a positive correlation between pupil size and spike rate. Across our population of recorded neurons, pupil size was also correlated with increases in the reliability of responses to sound, decreases in acoustic threshold, and no change in spectral bandwidth or best frequency. The changes in gain that we observed suggest that pupil size tracks the gross level of activity evoked by auditory stimuli: sounds become more salient to the rest of the brain when pupil is large. Changes in threshold and reliability may also support a more sensitive or precise representation of sound in high-arousal states.

### Stability of sensory tuning across pupil states

In mouse visual cortex, single neurons’ orientation selectivity increases when pupil is dilating rather than constricting: the neurons’ response to their preferred direction, but not the orthogonal direction, increases (Reimer et al., 2014). Although the impact of this change in orientation selectivity on tuning bandwidth has not been quantified, it suggests a narrowing of orientation tuning in the mean response across the population (Reimer et al., 2014). Our observation of stable tuning bandwidth in A1 may therefore indicate differences between how pupil-indexed states affect sensory selectivity in visual and auditory cortex.

Studies of task engagement effects on A1 reveal enhancement or suppression of neural responses to task-relevant sound features, including frequency (Fritz et al., 2003; David et al., 2012), amplitude modulation (Niwa et al., 2012), and spatial position (Lee and Middlebrooks, 2010), as well as generalized changes in excitability (Miller et al., 1972; Otazu et al., 2009). We found that best frequency is stable across large changes in pupil size, in contrast to previous studies of task-related plasticity in ferret primary auditory cortex that show frequency-specific effects (Fritz et al., 2003; David et al., 2012), suggesting a key difference between tasks that involve manipulation of the behavioral relevance of specific sound features and uncontrolled variation within passive states. Our data therefore suggest that effects of task engagement on frequency tuning in primary auditory cortex are not the result of non-specific increases in arousal, but instead involve separate mechanisms, such as feedback from frontal cortex (Fritz et al., 2010) or pairing activation of neuromodulatory systems involved in pupil dilation with specific auditory stimuli (Froemke et al., 2007; Martins and Froemke, 2015). The random sequences of tones used to characterize tuning would not pair release of acetylcholine with specific tone frequencies, and therefore would not be expected to gate shifts in tuning (Weinberger, 2004).

### Changes in neural excitability

A previous study in mouse auditory cortex found that intermediate pupil diameter was associated with maximum evoked responses to sound and minimal spontaneous activity (McGinley et al., 2015a). Although we observed some cells with non-monotonic relationships between pupil size and spike rate, this was the not the predominant pattern in our sample. In addition, we found that the effect of pupil-associated state usually had the same sign for both spontaneous and sound-evoked activity within a single neuron, in contrast to earlier results suggesting that intermediate pupil sizes were associated with opposite changes in spontaneous and evoked activity (McGinley et al., 2015a).

Several factors could explain this inconsistency between studies. The difference could reflect sampling of cortical layers. The earlier study targeted layers 4/5. We did not target any particular layer, but our method of recording auditory cells as the electrode advanced through cortex may have biased our sample towards more superficial layers. It is also possible that, compared to the earlier study, we tended to sample a different range of cortical states, either due to differences in experimental technique (the earlier study recorded head-restrained animals on a treadmill, while we did not use a treadmill) or differences in the behavioral patterns of ferrets and mice. Ferrets were not able to run and rarely showed substantial motor activity during the recordings. Thus, they may not have achieved the very high arousal state observed during bouts of running and other motor activity in mice.

### Pupil as an index of arousal or cognitive engagement

We used absolute pupil size as a measurement of cortical state. Our work therefore complements human pupillometry’s traditional focus on small, rapidly-decaying changes in pupil size that coincide with behavioral events (Kahneman and Beatty, 1966; Beatty, 1982a). The amplitude of these task-evoked dilations depends on a variety of cognitive variables (Zekveld et al., 2018). Although the variations in pupil size we observed provide a measurement of physiological arousal, they do not allow us to infer changes in a particular cognitive state, given that they occurred outside a controlled behavior. However, mechanisms like those we observed may underlie correlations between absolute pupil size and the efficiency of responses to sensory stimuli in some tasks (Beatty, 1982b; Murphy et al., 2011; van Kempen et al., 2019).

### Pupil as an index of noradrenergic tone

Activity in neuromodulatory centers, particularly the noradrenergic and cholinergic systems, has been proposed as a mechanism underlying correlations between pupil size and the level of neural activity in sensory cortex (McGinley et al., 2015b; Reimer et al., 2016). Evidence from multiple labs, species, and experimental techniques indicates that central release of noradrenaline from locus coeruleus is causally involved in pupil dilation and that noradrenaline release in sensory cortex accompanies pupil dilation (Aston-Jones and Cohen, 2005; Murphy et al., 2014; Joshi et al., 2016; Reimer et al., 2016; de Gee et al., 2017; Liu et al., 2017; Lovett-Barron et al., 2017; Larsen et al., 2018).

Despite the substantial evidence linking pupil size to noradrenergic tone, the gain increases we observed during pupil dilation do not directly match in vivo measurements of the effects of noradrenaline in auditory cortex. Iontophoresis of noradrenaline increases the signal-to-noise ratio of some auditory cortex neurons via a decrease in spontaneous, but not sound-evoked, activity (Foote et al., 1975; Manunta and Edeline, 1997, 1999). The gain increases we observe persist after subtracting mean spontaneous activity and are present when analyzing data from the evoked period alone. It is possible that iontophoresis does not replicate the spatial distribution or temporal dynamics of noradrenaline release in non-anesthetized animals. It is also possible that the gain increases we observed were a result of multiple modulatory systems acting on auditory cortex, both directly and indirectly. Activity in the cholinergic and dopaminergic systems are also correlated with changes in pupil size (Reimer et al., 2016; de Gee et al., 2017; Larsen et al., 2018). Given that pupil size is correlated with activity in multiple neuromodulatory systems, our failure to directly replicate results from spatially-localized, single-neuromodulator experiments is unsurprising, and indicates the complexity of neuromodulation under more natural conditions.

### Involvement of other cortical and subcortical brain regions

Some of the trial-to-trial variability we observed in auditory cortex may also be due to feedback from motor cortex (Schneider et al., 2014). Pupil size varies even when mice are still, and explains more variability in neural activity than locomotion (McGinley et al., 2015a). However, in non-anesthetized mice motor activities such as running, whisking, and licking a water reward are associated with pupil dilation (Lee and Margolis, 2016; Stringer et al., 2019). Detailed tracking of face and body movements explains more variability than pupil alone in mouse visual cortex (Stringer et al., 2019) and dorsal cortex (Musall et al., 2018). We did not track detailed motor behavior in the current study, and thus the extent to which motor activity predicts trial-to-trial variability in our data is not known. However, our results add to the evidence that pupil is one of several variables related to movement that can be used to predict cortical activity.

Pupil-related changes in neural activity are not restricted to the cortex. The same study that showed evidence of a causal relationship between pupil dilation and LC activity showed a similar relationship between pupil dilation and activity in superior and inferior colliculus (IC), including spiking preceding pupil dilation by hundreds of milliseconds and pupil dilation induced by microstimulation (Joshi et al., 2016). Some of the effects of arousal state we observed are likely to be inherited from auditory signals which must pass through IC before reaching cortex.

### Sleep states during head-restrained recordings

Recent results in head-restrained mice showed fluctuations between awake, rapid eye-movement (REM) sleep, and non-REM sleep states associated with changes in pupil size, with constricted pupil associated with REM sleep (Yuzgec et al., 2018). We observed epochs of tonically constricted pupil accompanied by an abrupt increase in saccade rate. Although alpha power consistently increased as pupil constricted, this trend reversed itself during these epochs. We speculate that over the course of lengthy, head-restrained passive recordings, the ferrets may become progressively more drowsy, and that the high-saccade, low-alpha epochs indicate sleep onset.

The pattern of constricted pupil, saccades, and lack of eyelid closure we observed may also be an artifact of head-restrained recording. Methods for recording the activity of single neurons in freely-moving animals can be supplemented with head-mounted cameras to track pupil (Meyer et al., 2018). It is likely that these techniques will yield data on changes in internal state more relevant to natural environments.

Our observation of putative sleep states, like previous data from mice (Yuzgec et al., 2018), complicates the suggestion that the pupil size of head-restrained animals reveals a continuum of awake arousal states analogous to, but distinct from, sub-states of sleep (McGinley et al., 2015b). Thus, a full characterization of arousal states must incorporate both pupillometry and other physiological measurements that can distinguish between drowsy waking states and sleep.

## Acknowledgments

This work was supported by grants from the National Institute on Deafness and Other Communication Disorders (R01 DC0495, R21 DC016969, F31 DC016204) and the ARCS Foundation Oregon Chapter. The authors would like to thank Daniela Saderi, Luke A. Shaheen, and Sean Slee for training, advice, and assistance with neurophysiological recording, Ulysses Duckler for animal care, and Matthew J. McGinley, Avinash Bala, and Robert Peterka for technical assistance with pupillometry. The authors declare no competing financial interests.

## Supplementary Material

**Movie 1.**
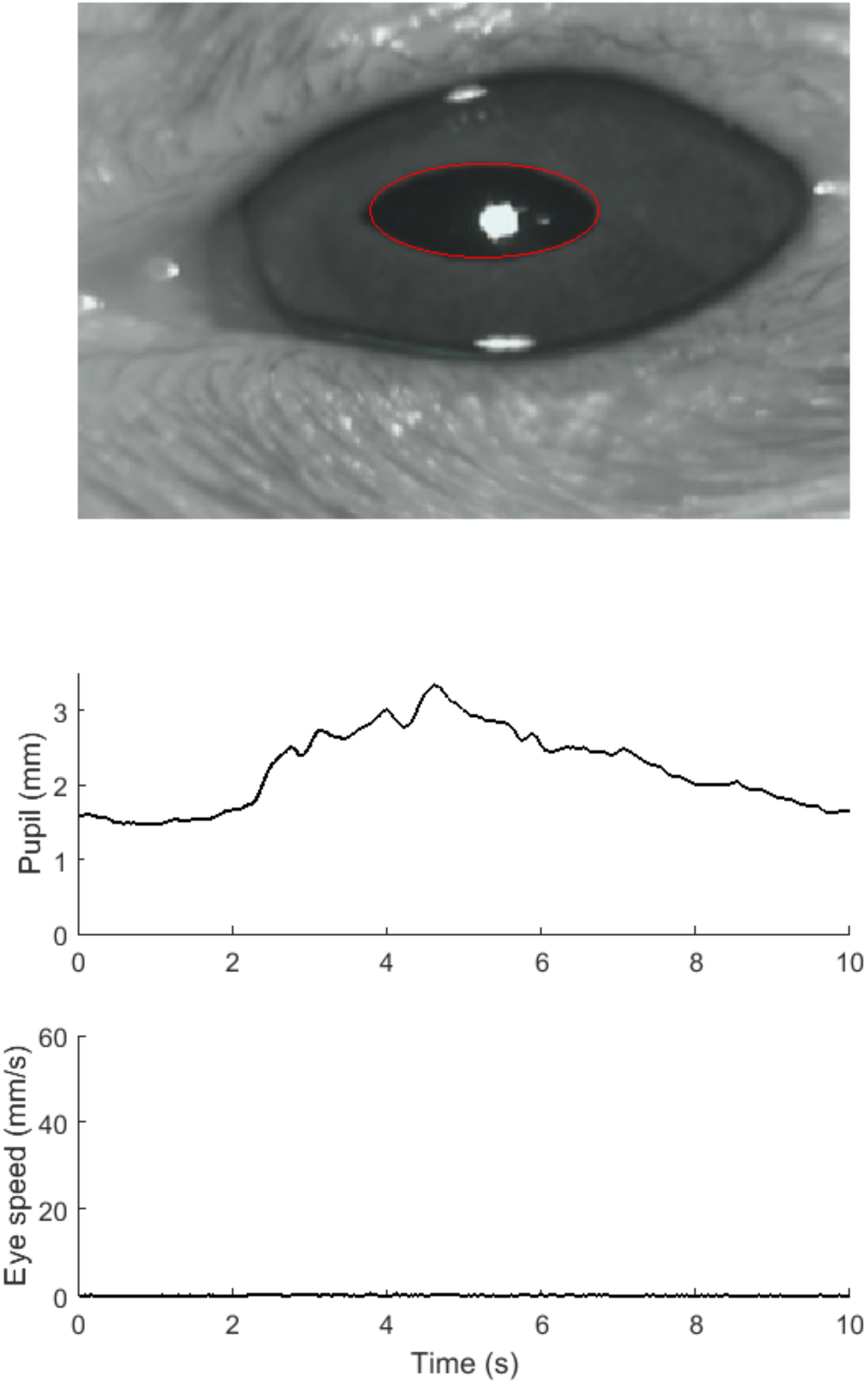
Real-time example of pupil size and eye movement dynamics during passive, awake state.

**Movie 2.**
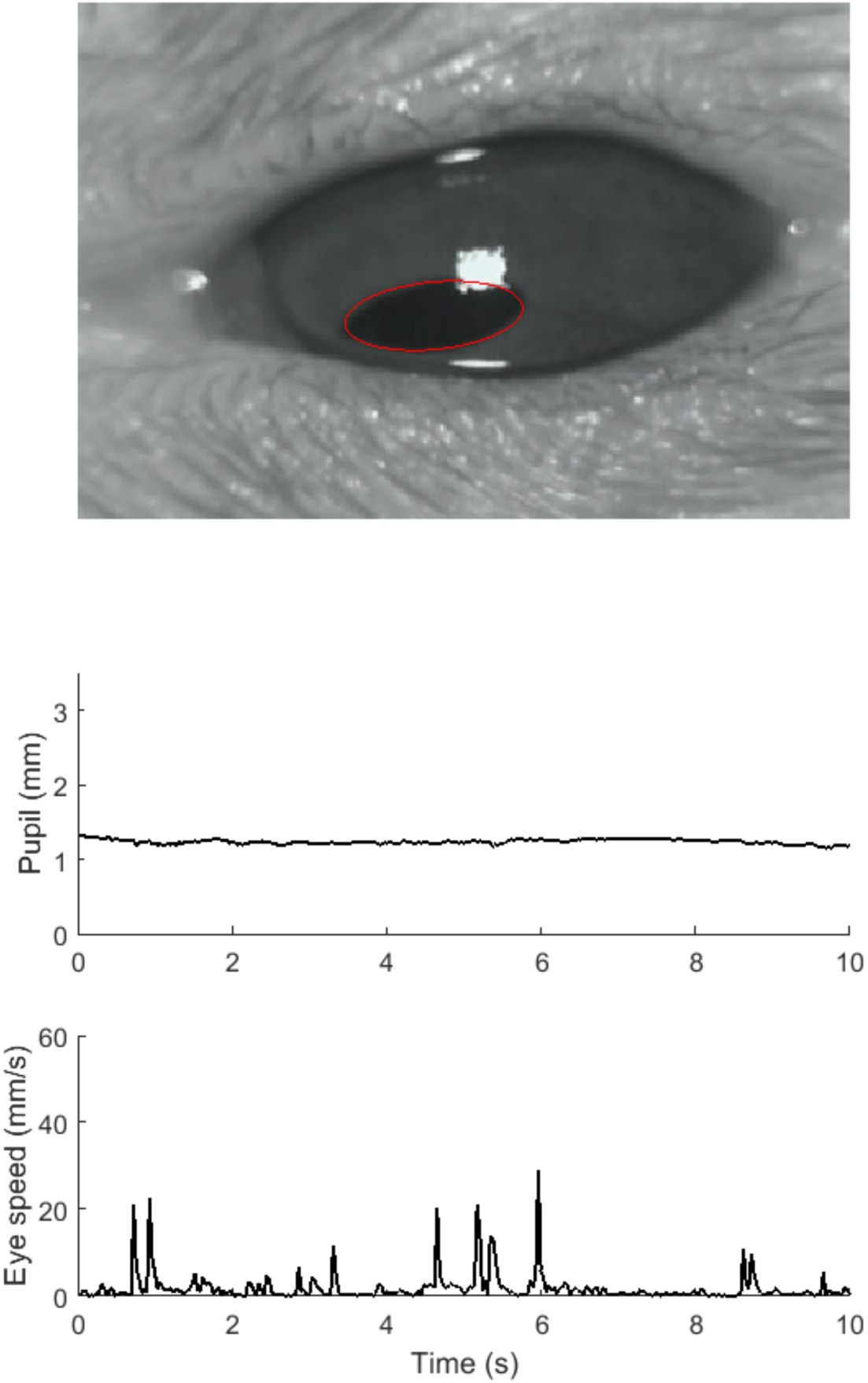
Real-time example of pupil size and eye movement dynamics during sleep state.

## Footnotes

1 Since the expected value of the Gamma distribution is α/β, we can compute the posterior for the average spontaneous rate of the population as *b_α_*/*b_β_* when the pupil is small and *b_α_*/*b_β_*× *bg_μ_* when the pupil is large.

